# Layered phage therapy exploits capsule-loss escape and altered antibiotic susceptibility in CG23 hypervirulent *Klebsiella pneumoniae*

**DOI:** 10.64898/2026.10.20.760367

**Authors:** Ayuni Norman, Jun Hao Liew, Lalit Mohan, Chanvutha Ron, Aryeh Chiam, Yan Sun, Indira Kartigayen, May Delos Santos, Yingying Li, Bernice Siu Yan Tan, Shu Xuan Tan, Raksha Anand, Evangeline, Kay En Low, Isabelle Bonne, Andrea Lay Hoon Kwa, Barry Kreiswirth, Yunn-Hwen Gan, Roland Huber, Pablo Bifani

## Abstract

The rise of drug-resistant *Klebsiella pneumoniae* infections is prompting a re-evaluation of phage therapy as an alternative treatment. Phage therapy has achieved successful outcomes, yet important limitations persist, including the rapid emergence of phage resistance and the time required to source suitable phages. Combining multiple phages into a phage cocktail based on genetic diversity, morphology or distinct infection mechanisms is a common strategy to mitigate resistance. Phage pairing based on these criteria neither mitigates nor delays the emergence of resistance, using CG23 hypervirulent *Klebsiella pneumoniae* (hv*Kp*) EC0675 as a model. In CG23 *Kp* strains, resistance arises through disruption of the capsular biosynthetic pathway, leading to capsule impairment and global cross-resistance to all capsule-active phages. Combining phages active against encapsulated wild-type bacteria with those active against capsule-deficient escape mutants can overcome this constraint, improving bacterial suppression *in vitro* and reducing bacterial burdens across multiple organs in a mouse systemic infection model. In parallel, phage-driven capsule loss increased susceptibility of hv*Kp* EC0675 to selected antibiotics targeting the bacterial envelope or cell wall *in vitro*. These findings highlight the contribution of capsule-associated resistance to phage cross-resistance in the CG23 *Kp* strains examined, and support further evaluation of phage combinations targeting both wild-type bacteria and resistant derivatives, including their use alongside selected antibiotics, informing future clinical translation.

## Introduction

Phage therapy has undergone a significant revival in response to the mounting challenge of antibiotic resistance, a development fuelled by urgent clinical needs and transformative advances in phage biology, genetic engineering, and AI-based optimization^1^ as evidenced by the registration of at least 32 clinical trials worldwide between 2020 and 2024^2^. The use of phage therapy to treat *Klebsiella pneumoniae* (*Kp*), an ESKAPE organism on the WHO priority list, is no exception, with 13 reports describing its application up to 2025 (Supplementary Table 1). *Kp* is a gram-negative opportunistic pathogen found ubiquitously in the environment, as well as contaminants on medical devices^3^. It carries various virulence genes and acquires antibiotic resistance, including extended-spectrum β-lactamases (ESBLs)^4^. It can successfully evade the host immune system due to its capsule, which contributes to its spread and the formation of biofilms^5,6^, and can colonize the mucosal surfaces (the upper respiratory and gastrointestinal tract). Not surprisingly, it is a leading cause of nosocomial pneumonia^5,7^.

*Kp* has been grouped into two pathotypes: classical (c*Kp*) and hypervirulent (hv*Kp*). c*Kp* is primarily associated with nosocomial infections in immunocompromised patients and tends to be multidrug-resistant^8,9^. The term hv*Kp* has been used loosely to refer to hypervirulent or hypermucoviscous *Kp,* associated with community-acquired infections in mainly healthy young and immunocompromised hosts^10^. However, it has also been noted that not all hypermucoviscous *Kp* are hypervirulent^11^. Both c*Kp* and hv*Kp* can cause infections, including pneumonia, liver abscess, meningitis, septic arthritis, and urinary tract infections^4,12^. Severe infections caused by hv*Kp* include pyogenic liver abscesses, bacteremia, meningitis, endophthalmitis, and necrotising fasciitis^13^. Recently, many countries have reported sporadic emergence and mortality due to convergent multidrug-resistant and hypervirulent *Kp* (MDR-hv*Kp*) strains and pan-drug-resistant (PDR) hv*Kp* strains^4,10,14,15^. Carbapenems, considered last-resort antibiotics, are becoming less reliable due to increasing resistance^16^. Moreover, recent findings from Asia, specifically China, indicate that carbapenem-resistant hv*Kp* poses a growing health threat due to its heightened virulence coupled with escalating resistance. Specific sequence types (STs), including ST11, ST15, and ST23, have been identified as predominant, with ST23 exhibiting notably enhanced virulence^17^.

Growing concerns about multidrug resistance, along with advances in understanding phage biology, have increased interest in bacteriophages as potential therapeutic alternatives. Previous research involving phage-resistant mutants and phage–antibiotic combinations suggests that these approaches may help identify treatments active against both parental bacterial populations and their resistant derivatives^18–22^. However, effectiveness may depend on the bacterial background and experimental context, as resistance mechanisms, phage host ranges, and antibiotic responses can vary across strains and conditions. Here, we developed phage cocktails against hv*Kp*, where we chose to target clonal group 23 (CG23) hv*Kp* as a model, given its prevalence and relevance in the Asia-Pacific region, particularly in association with pyogenic liver abscesses^23,24^. CG23, comprising ST23 and related sequence types including ST26, ST57, and ST163^25^ is a major hypervirulence-associated lineage of *Kp* in which multidrug-resistant strains, including carbapenemase-producing strains, have emerged^26–29^. Our layered phage cocktail, designed by pairing phages targeting wild-type and phage-resistant mutants, counteracts phage resistance and could yield an effective combination against hv*Kp*, with or without antibiotics.

## Results

Early into this study, we found that *Kp* phages have a propensity to be highly strain- or sequence-type-specific, as per other studies^30,31^. Consequently, we focused on CG23, a major hypervirulence-associated *Kp* lineage^29^, given its predominance in Asia and its association with pyogenic liver abscesses, and confirmed that the wild-type isolates examined in the string-test experiments exhibited a hypermucoviscous phenotype^17,32^. This group serves as a model for developing ready-to-use, pre-optimised phage cocktails that can be cleared for clinical use. In **Figure 1a**, we summarised our workflow, outlining the effective phage pairing approach (pairing phages targeting wild-type and phage-resistant populations) to overcome bacterial resistance during treatment of encapsulated wild-type *Kp* in both *in vitro* and *in vivo* settings, with and without antibiotics.

**Fig. 1:**
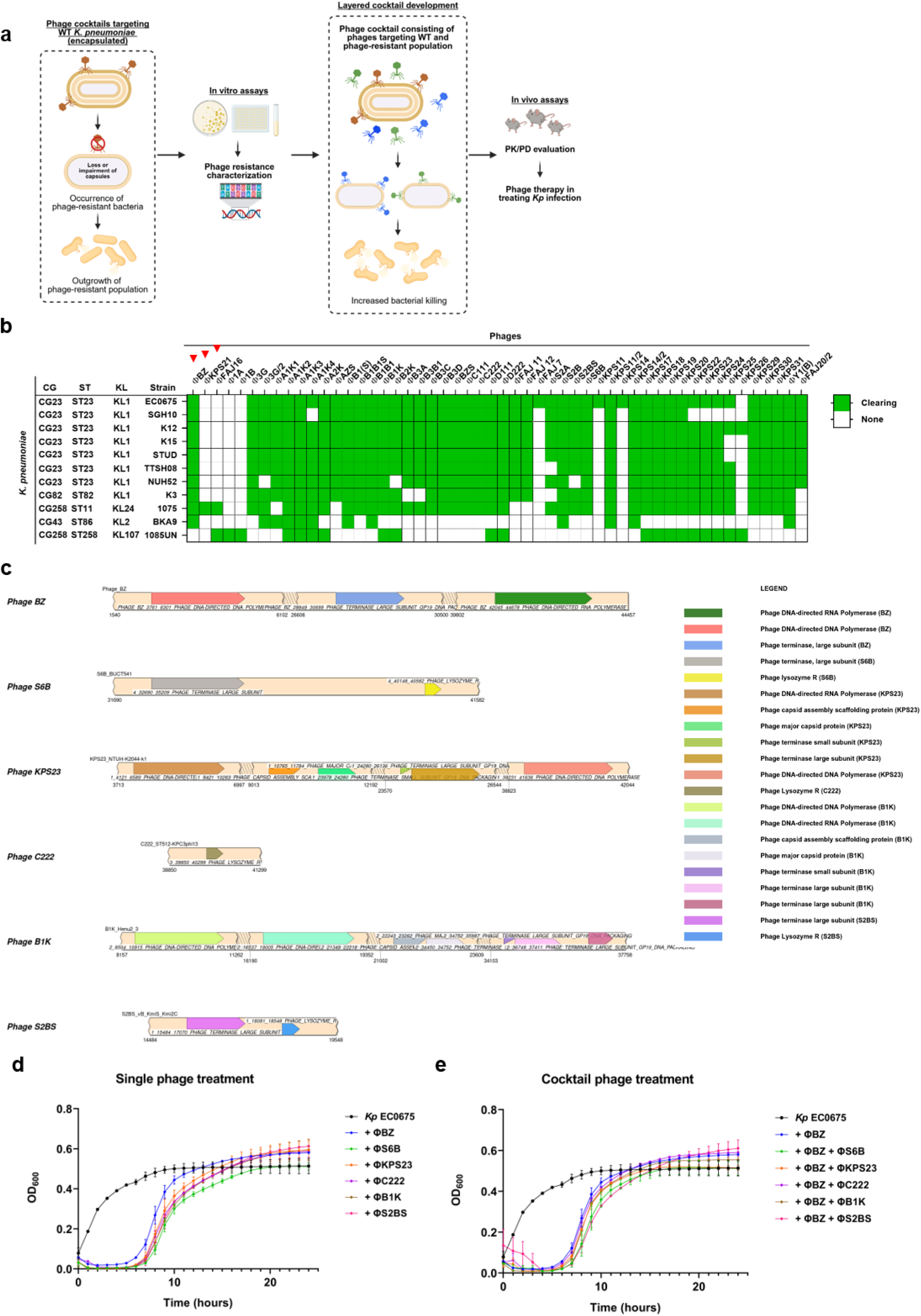
Diverse phage combinations tested on hypermucoviscous *K. pneumoniae*. **(a)** Workflow of phage pairing approach to design an effective phage cocktail for therapeutic use. **(b)** Representative phage host susceptibility against various *Kp* strains. ST: Sequence type, KL: K locus. Red arrows indicate the main phages focused on in this study. **(c)** SimpleSynteny webserver was used to visualise and compare synteny across the phage genomes. Comparison of annotated phage genes in selected CG23-targeting phages reveals their diversity. **(d-e)** The bacterial killing curve of single (d) and cocktail (e) phage treatments against *Kp* EC0675 strain (all data shown at 1h interval readings at OD_600_ represent three independent experiments, bars indicate mean ± SEM).

A total of 147 phages were isolated from environmental and wastewater samples collected in Singapore. Various sequence types (ST) within 10 CGs were used as hosts for phage isolation. The isolated phages were confirmed for specificity against CG23 isolates, and validated on a well-characterised and representative collection of clinical *Kp* isolates from the USA and Singapore (**Extended Data Fig. 1a**). Selected phages that target *Kp* CG23 strains are shown in **Fig. 1b** and **Extended Data Table 1**.

### Phage cocktails targeting wild-type CG23 strains do not mitigate the emergence of phage resistance

Six phages (ΦBZ, ΦS6B, ΦKPS23, ΦC222, ΦB1K, and ΦS2BS) were selected based on their broad host range and lytic potency in spot assays and subsequently evaluated in kill-kinetic assays against the model strain *Kp* EC0675. Genomic comparison showed substantial diversity, although these are all CG23-targeting phages **(Fig. 1c**). In all cases, an initial inhibition of bacterial growth was observed until approximately 5 to 9h post-infection (**Fig. 1d**). Thereafter, steady bacterial growth was observed with all the phages. Consequently, phage cocktails were prepared to mitigate bacterial growth (**Fig. 1e**), yet these cocktails did not delay the emergence of resistance, in contrast to other studies^33–35^. The effect of phage combination was neither synergistic nor additive, as often seen for disinfectants or antimicrobials^36^.

### Loss or impairment of capsules in CG23-resistant mutants

We selected ΦBZ, a potent and broad host-range phage, as a model for this study. Genomic analysis of ΦBZ showed a genome size of 45.4 kbp with a GC content of 45.95% (**Fig. 2a**), and BLAST genome analysis revealed its sequence similarity at 99.19% to *Klebsiella* phage Roth32 (7801.1) with a query coverage of 99%. ΦBZ was experimentally determined to be a lytic phage, supported by sequencing data. No virulent and toxic sequences (e.g., toxins or antibiotic-resistance genes) were detected. Electron microscopy demonstrated that ΦBZ exhibits a morphology characterized by an isometric head and a long, flexible, noncontractile tail, consistent with the siphovirus morphotype (**Fig. 2b**). *Kp* EC0675 was treated with ΦBZ, and phage-resistant mutants were isolated to investigate the mechanisms underlying phage resistance and hypermucoviscosity (**Fig. 2c-e)**. The phage-resistant isolates of EC0675 exhibited distinct phenotypes: smaller, varied in size, and appeared translucent, indicating a loss or diminished capsule production (**Fig. 2d**). The phage-resistant isolates were also observed to have decreased mucoviscosity compared to the wild-type (**Fig. 2c-e and Extended Data Fig. 1b**). Additionally, eight CG23 strains were also treated with ΦBZ, and wild-type bacteria and phage-resistant mutant counterparts were assessed with the string test. All phage-resistant mutants were non-hypermucoviscous, whereas all wild-type isolates showed hypermucoviscosity (P<0.001) (**Fig. 2e**). Mutation analysis of seven randomly selected ΦBZ-resistant isolates (ΦBZ-RI 1-7) revealed mutations in the genes associated with the biosynthesis of capsular polysaccharides (*wcaJ*, *wzb,* and *wzc* genes), consistent with their impaired capsule phenotype (**Fig. 3a**). Most resistant mutants encoded alterations in one of the coding sequences of the capsular biosynthetic genes, underscoring their impaired capsule production **(Fig. 3b)**. This impaired capsule production among the mutants was responsible for the curtailed adsorption of ΦBZ when compared to their wild-type counterpart (**Fig. 3c**). The disparity in capsule production between mutants and wild-type was determined using quantitative measurement of uronic acids, a major capsular polysaccharide^37,38^. The uronic acid content of the ΦBZ-resistant mutants was significantly lower than that of wild-type *Kp* EC0675 (**Fig. 3d**).

**Fig. 2:**
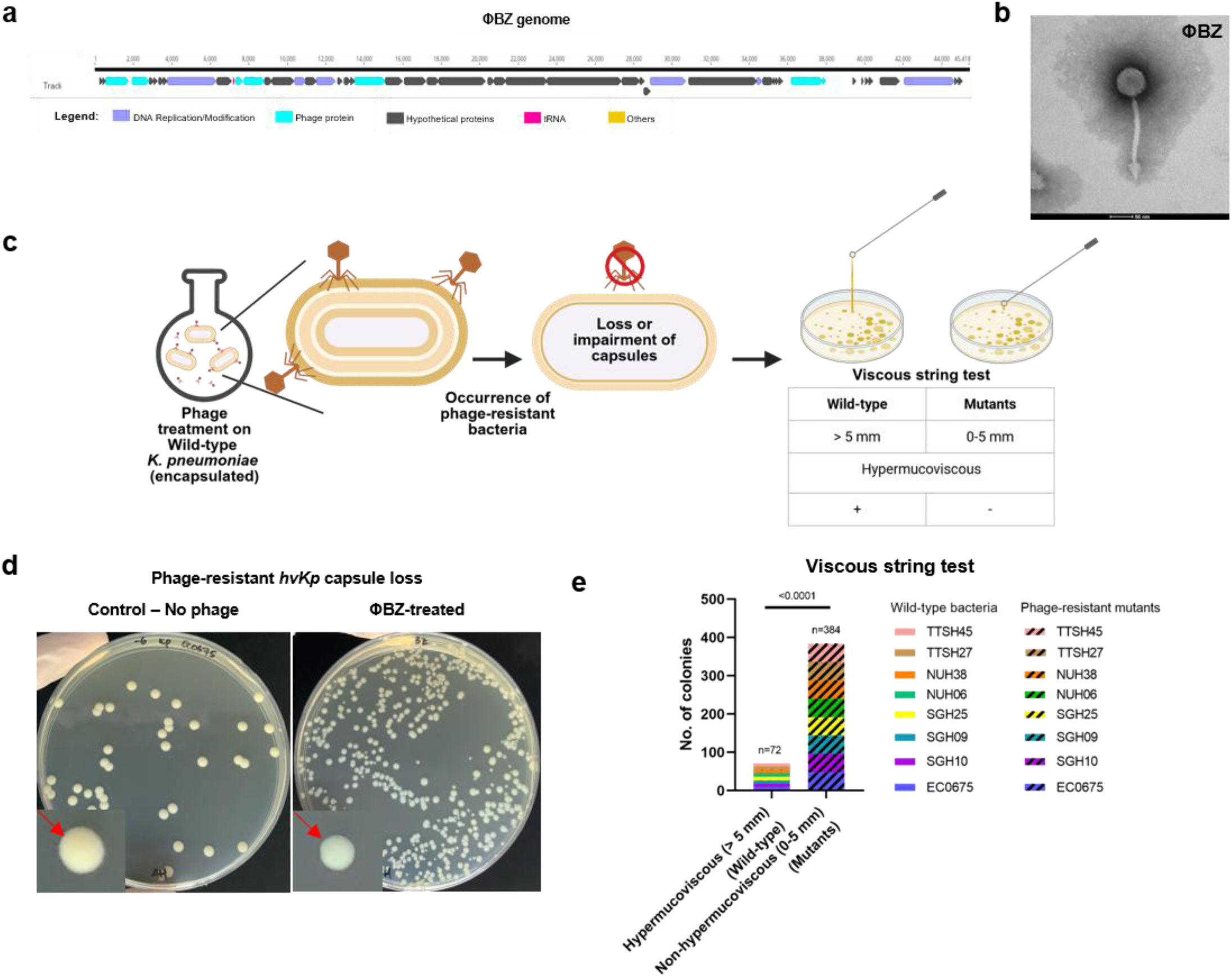
Characterisation of primary phage ΦBZ and phage-resistant mutants across clinical CG23 *Kp* strains. **(a)** Genome map illustration of ΦBZ. **(b)** TEM representative image of ΦBZ. **(c)** Schematic summary of phage resistance assay. Phage treatment of wild-type bacteria led to the occurrence of phage-resistant bacteria that had lost or impaired capsules. Viscous string tests highlighted hypermucoviscous-positive for wild-type bacterial strains and hypermucoviscous-negative for the phage-resistant mutants. **(d)** Resistance emerging upon treatment of ΦBZ with *Kp* EC0675 strain (data shown are representative of 4 independent experiments), morphological difference of bacterial colonies in *Kp* EC0675 cultures of control (no phage treatment) and ΦBZ-treated after 24h in mixture. **(e)** Viscous string tests were conducted for 8 clinical wild-type bacterial strains (n=72 isolates) and their phage-resistant mutants (n=384 isolates). All wild-type isolates were tested hypermucoviscous-positive, and all phage-resistant mutants were tested hypermucoviscous-negative. Fisher’s exact test (two-tailed) was conducted to analyse the string test hypermucoviscosity results comparing wild-type and mutant isolates.

**Fig. 3:**
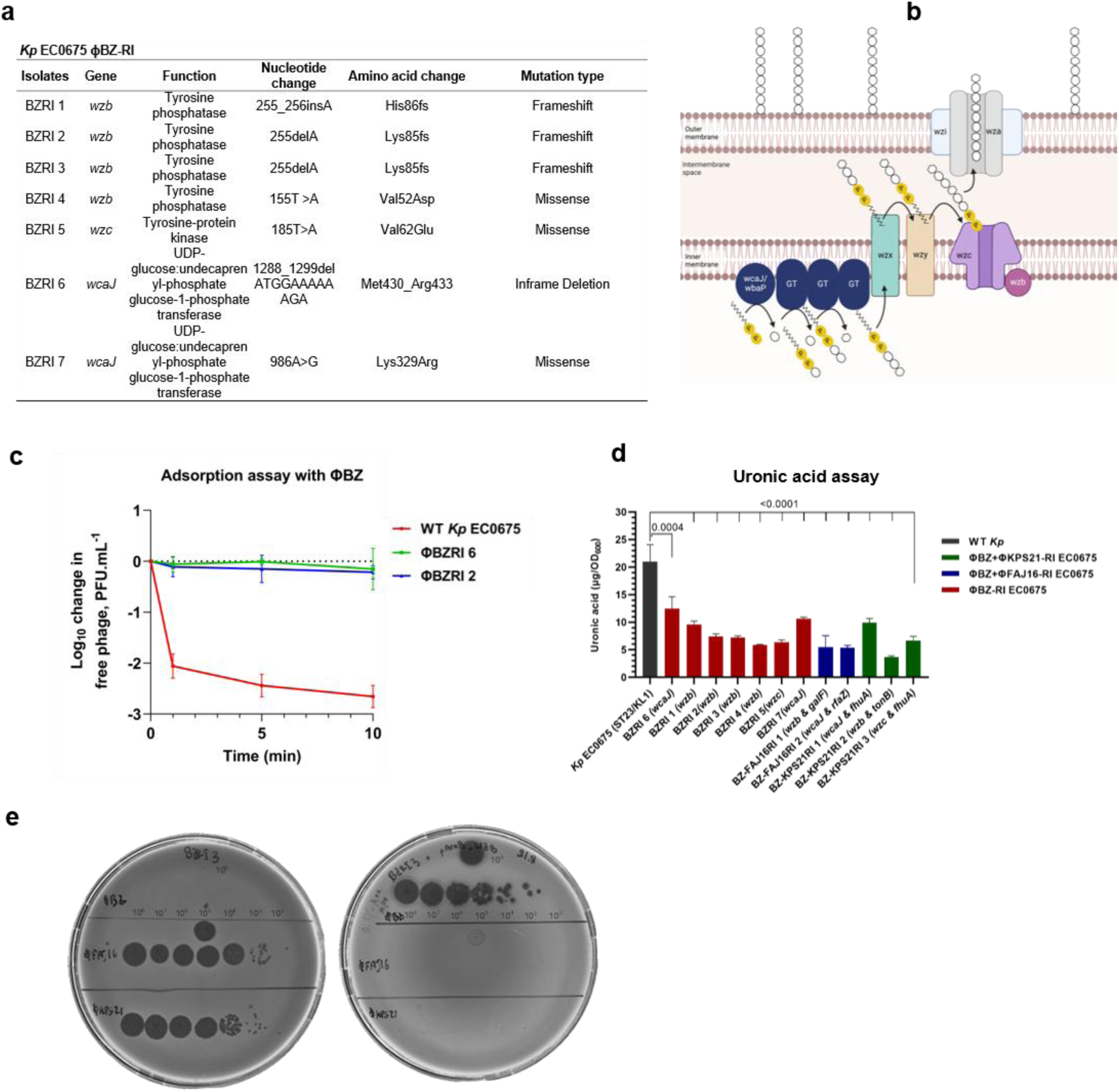
Genotypic and phenotypic characterisation of ΦBZ phage-resistant mutants. **(a)** Mutation analysis of ΦBZ-resistant isolates (ΦBZ-RI). **(b)** Schematic diagram of capsule biosynthesis pathway in *Kp*. **(c)** Adsorption assay of *Kp* EC0675 WT and ΦBZ-RIs with ΦBZ to determine whether impaired capsule production led to the loss of phage adsorption. n=3 biological replicates; bars indicate mean ± SD. **(d)** Capsule production quantitation via uronic acid content of cultures grown to OD_600_ 1.0 to determine the effect of mutation in the EC0675 ΦBZ-RIs compared to the EC0675 WT. Gene mutations for each ΦBZ-RI are labelled in brackets after its isolate name. n=3 biological replicates; bars indicate mean ± SD. One-way ANOVA tests with multiple comparisons were performed by comparing the mean of each biological triplicate of ΦBZ-RI with *Kp* EC0675 WT. (e) Complementation of the ΦBZRI 3-resistant mutant with wild-type *wzb* restored susceptibility to ΦBZ. Left plate: ΦBZRI 3 and right plate: ΦBZRI 3 complemented with wild-type *wzb*.

### Complementation of ΦBZRI 3 with wild-type *wzb* re-sensitized to ΦBZ

To confirm that the loss of capsule by *wzb* mutation led to resistance to ΦBZ, we cloned the *wzb* gene from wild-type *Kp* EC0675 strain into vector pAraB-Apr. The complemented ΦBZRI 3 was screened for phage susceptibility, and susceptibility to ΦBZ was restored (**Fig**. **3e**). Upon complementation, the mutant lost its susceptibility to ΦKPS21 and ΦFAJ16, behaving like the wild-type *Kp* EC0675 strain.

### Validation of loss or impairment of capsules across other CG23 isolates and CG23-specific phages

The loss or impaired production of the capsule associated with phage resistance was not exclusive to ΦBZ-EC0675 pairing. Phage-resistant EC0675 mutants and mutants from another *Kp* SGH10 clinical strain, when subjected to selective pressure with six randomly selected CG23-specific diverse phages (ΦBZ, ΦS6B, ΦKPS23, ΦC222, ΦB1K, and ΦS2BS), also exhibited impaired capsule production in all instances (**Extended Data Fig. 2a-b**). Likewise, similar results were observed for eight random pairings of ΦBZ to CG23 strains **(Extended Data Fig. 2c)**. Phage selection for this experiment was based on genetic diversity (**Fig. 1c**) to emphasize that the mechanism of resistance is global and not associated with any specific phage genotype or phage-bacteria pairing. Note that specific phage receptors have not been determined for any of the *Kp* phages.

### Phage resistance in hv-*K. pneumoniae* is global

While phage combinations targeting the hv*Kp* did not mitigate or delay the emergence of phage resistance; here we additionally observed that resistance to one phage can confer either full or partial resistance to all *Kp* phages tested (**Fig. 4a**), encompassing all clinical isolates as shown in **Fig. 2e**. ΦBZ-resistant mutants while gaining resistance to ΦBZ (*wzb*, *wzc* and *wcaJ*) also exhibited partial or complete resistance to all phages isolated using CG23 *Kp* strains (**Fig. 4a**). Overall, this data suggests ΦBZ and other CG23 phages are likely capsule dependence for infection, requiring a highly expressed capsule for adsorption and subsequent infection. The impairment of capsular biosynthesis suggests its possible correlation to the observed phenotypic changes in the phage-resistant isolates. The cross-resistance associated with the diminished capsule likely explains the lack of synergistic or additive phenotypes in phage cocktails targeting the wild-type hv*Kp*.

**Fig. 4:**
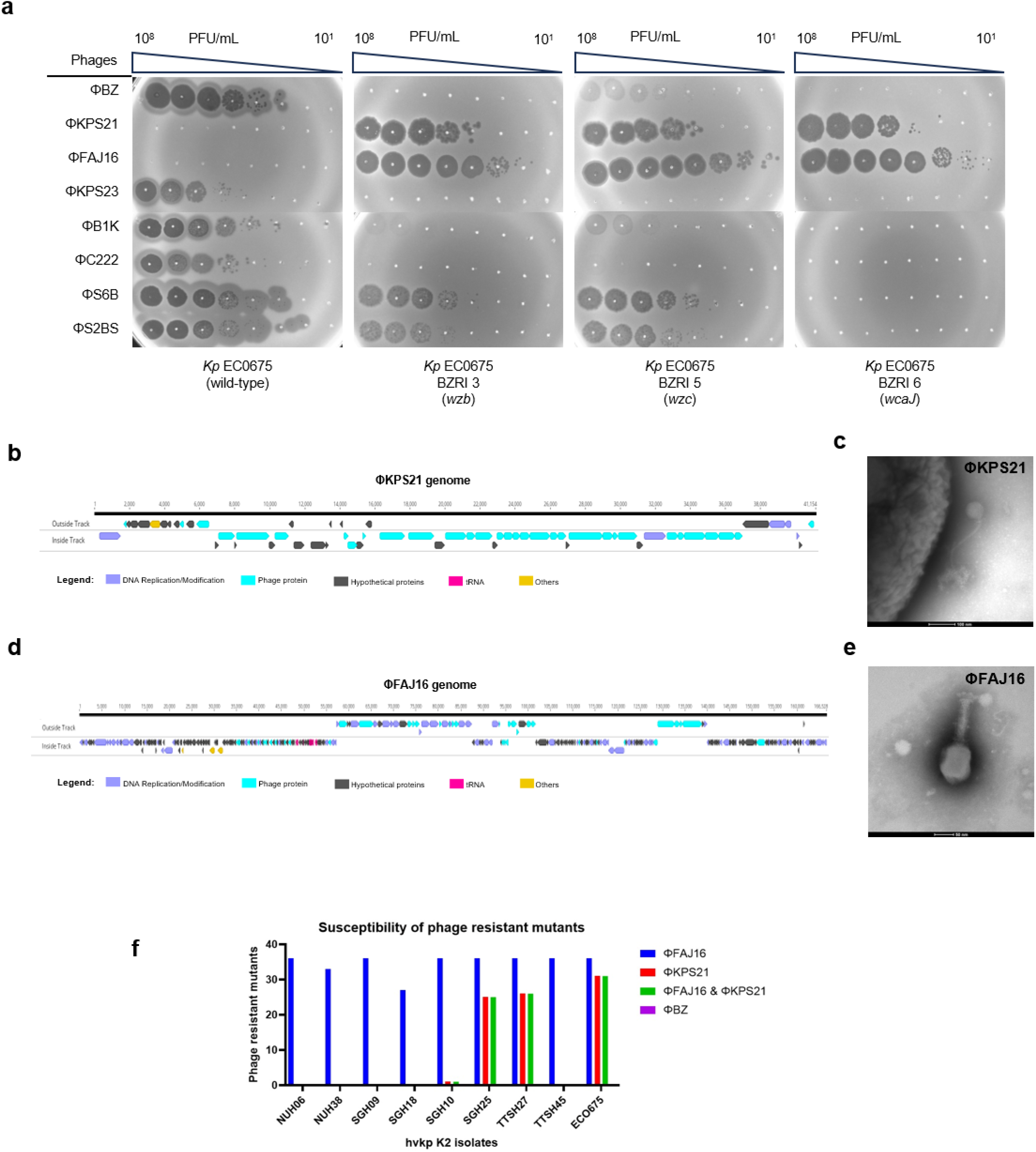
Characterisation of second-layer phages targeting phage-resistant mutants. **(a)** Phage host susceptibility against *Kp* EC0675 and ΦBZ-RI mutants. CG23-specific phages: ΦBZ, ΦKPS23, ΦB1K, ΦC222, ΦS6B and ΦS2BS, and selected second-layer phages: ΦFAJ16 and ΦKPS23. **(b)** Genome map illustration of ΦKPS21. **(c)** Representative TEM image of ΦKPS21. **(d)** Genome map illustration of ΦFAJ16. **(e)** Representative TEM image of ΦFAJ16. **(f)** Phage susceptibility profile of hv*Kp* ΦBZ-resistant mutants of various clinical isolates. ΦFAJ16, ΦKPS21 and both (n=36 isolates per strain). No phage susceptibility were observed for ΦBZ in these mutants.

### Identification and characterisation of phages targeting impaired capsular mutants

ΦBZ and its combinations (**Fig. 1e**) did not mitigate the emergence of phage resistance, necessitating a re-strategized cocktail development towards a layered approach. We screened ΦBZ-resistant mutants with impaired capsule production for new active phages. Among the 98 phages evaluated from our collection, ΦKPS21 and ΦFAJ16 emerged as the most promising candidates, demonstrating efficacy against all ΦBZ-resistant mutants in terms of potency and strain broadness (**Fig. 4a**). ΦKPS21 and ΦFAJ16 were used to design a layered phage cocktail and to generate secondary phage mutants.

ΦKPS21 and ΦFAJ16 were characterised phenotypically and genotypically (**Fig. 4b-e).** Although genomic analysis of ΦKPS21 identified an integrase and repressor gene consistent with a temperate lifestyle, ΦKPS21 demonstrated clear lytic activity against capsule-deficient EC0675 mutants under the experimental conditions tested (**Fig. 4a**). For the purposes of this study, ΦKPS21 served as a mechanistic tool to characterise secondary resistance pathways and validate the layered cocktail concept; its inclusion does not imply suitability for direct clinical application. Genomic analysis of ΦKPS21 and ΦFAJ16 revealed genome sizes of 41.1 kbp with a GC content of 50.67% and 166.5 kbp with a GC content of 39.60%, respectively. ΦFAJ16 was experimentally determined to be a lytic phage, supported by sequencing data. No virulent and toxic sequences (e.g., toxins or antibiotic-resistance genes) were detected for both ΦKPS21 and ΦFAJ16. Electron microscopy revealed that ΦKPS21 and ΦFAJ16 resemble members of siphovirus and myovirus morphotypes, respectively. Additionally, resistant mutants from eight other CG23 isolates were generated against ΦBZ and were tested for their sensitivity against ΦFAJ16 and ΦKPS21(**Fig. 4f).** The resistant mutants from the other CG23 isolates were also susceptible to either ΦFAJ16, ΦKPS21 or both.

### Characterisation of secondary phage-resistant mutants generated from a layered phage cocktail

Primary ΦBZ-resistant mutants were used to generate the secondary phage-resistant colonies upon treatment with ΦKPS21. Mutation analysis of the three EC0675 ΦBZ+ΦKPS21-resistant mutants revealed frameshift mutations in the TonB subunit and ferrichrome outer membrane receptor, which may be potential target receptors of the phage (**Fig. 5a and 5b**). Notably, a frameshift mutation in TonB, observed in phage-resistant isolate BZ-KPS21RI 2, has also been previously reported^39,40^. Likewise, primary ΦBZ-resistant mutants were used to select for secondary resistant mutants to ΦFAJ16. Analysis of the ΦBZ+ΦFAJ16-resistant isolate reveals that all mutations were associated with the 5’ conserved region in the CPS locus. The mutant BZ-FAJ16RI 1 exhibits two mutations, one occurs in *wzb* due to exposure to ΦBZ, which leads to an impaired capsule chain length, as this gene is known to regulate CPS length^41^. This may lead to the exposure of receptors to which ΦFAJ16 might attach. The second mutation upon exposure to ΦFAJ16 was observed in *galF*, which plays a regulatory role in CPS biosynthesis^42,43^ (**Fig. 5c-e**). Its putative association with phage resistance has yet to be confirmed. Mutant BZ-FAJ16RI 2 has a primary mutation (caused by ΦBZ) in *wcaJ*, a glycosyltransferase responsible for the initiation of colanic acid synthesis and thus leading to impaired capsular synthesis^44^. The secondary mutation (caused upon exposure to ΦFAJ16) was observed in *rfaZ*, encoding a liposaccharide biosynthesis core protein^45^(**Fig. 5a**). Importantly, all the isolates of ΦFAJ16 and ΦKPS21-resistant mutants maintained the mutations originally present in the primary ΦBZ-resistant mutants (**Fig. 5a and 5c**).

**Fig. 5:**
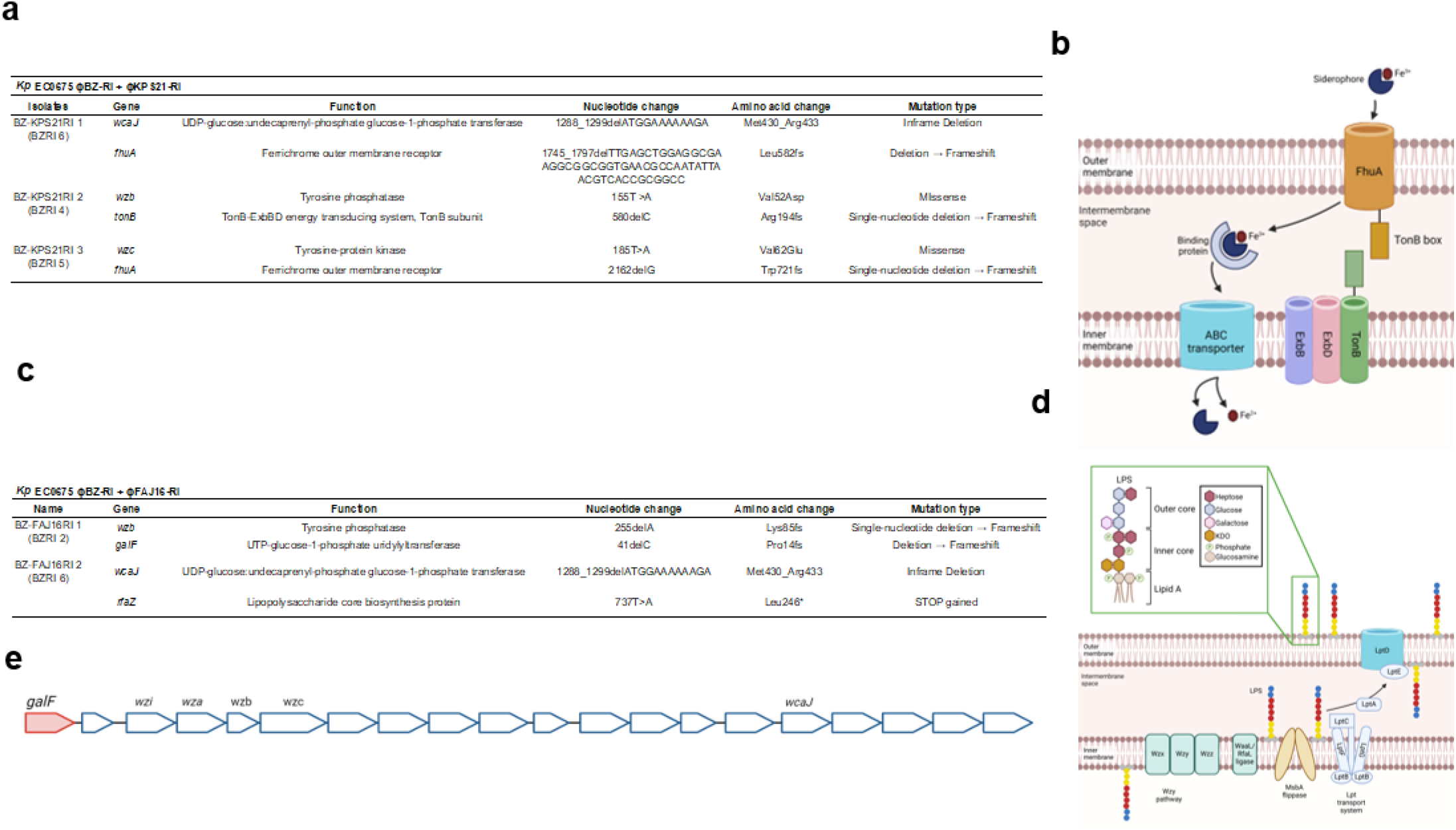
Mutation analysis of phage-resistant mutants generated from second-layer phages. **(a)** Mutation analysis of ΦKPS21-resistant isolates (ΦKPS21-RI) using ΦBZ-RI. **(b)** Schematic diagram of TonB-dependent transporters in *Kp*. **(c)** Mutation analysis of ΦFAJ16-resistant isolates (ΦFAJ16-RI) using ΦBZ-RI. **(d)** Schematic diagram of the lipopolysaccharide biosynthesis pathway in *Kp*. **(e)** Genetic loci illustration of K1 capsule in *Kp*. Arrows represent the different genes, *galF* in red corresponds to the region involved in the mutation of ΦFAJ16-RI 2. Conserved genes are labeled. These genes *wza*, *wzb*, *wzc,* and *wzi are* involved in the assembly and export of the capsule, while *wcaJ* is involved in initiating glycosyl-transferase.

### A layered phage cocktail enhances bacterial killing efficiency

The bacterial killing assay was designed to determine the efficiency of the layered treatment approach (**Fig. 6a**). Different ratios of ΦBZ, ΦFAJ16, and ΦKPS21 and bacteria demonstrated a range of killing efficiency at the various time points (**Extended data Fig. 3a**). Both ΦFAJ16 and ΦKPS21 were included as negative controls, as they do not target the wild-type EC0675 and only infect the emerging phage-resistant population. At 4h, similar effectiveness in controlling bacterial growth was observed for single- and multi-phage treatments. However, after 24h, the three-phage-cocktail treatment emerged as the most optimal combination compared to single- and two-phage-cocktail treatments **(Extended data Fig. 3a)**. The introduction of a second layer of phages that targets the phage-resistant wild-type population was able to reduce the overall bacterial load. Similarly observed in the kill kinetic assay, the phage cocktail setups portrayed superior killing activity, with a delayed emergence of phage resistance (∼8 to 10h) and lower bacterial density at 24h compared to the single ΦBZ treatment (**Fig. 5b**). Growth rate constants (K) were determined for the various phage treatments setups (**Fig. 5c**). Similar growth rates were observed for the control group with no phage treatment (K=0.1739) and the single-phage treatments that do not target the wild-type EC0675 (K=0.1636 for ΦKPS21 and K=0.1678 for ΦFAJ16). The single-phage ΦBZ treatment exhibited a lower growth rate (K=0.1355), while two- and three-phage treatments demonstrated even lower growth rates (K=0.0844 for ΦBZ+ΦKPS2, K=0.0996 for ΦBZ+ΦFAJ16, and K=0.0902 for the three-phage treatment), highlighting the bacteria’s limited ability to thrive under phage treatments and even more limited under phage combination treatments. The *in vitro* killing activity of the phages against EC0675 was further assessed by CFU enumeration (**Fig. 5d**). At 25 minutes post-treatment, single-, two-, and three-phage treatments significantly reduced bacterial counts compared to the control (*p* <0.01). At 4h, both single- and two-phage treatments continued to show reduced bacterial loads relative to the control (*p* <0.01). However, after 24h, the emergence of phage-resistant mutants resulted in a higher bacterial load in the single-phage ΦBZ treatment compared to other treatments. Moreover, all phage treatment groups displayed reduced bacterial loads, with the pairing of ΦBZ and ΦKPS21 resulting in lower bacterial counts than the pairing of ΦBZ and ΦFAJ16 when compared to the control. Notably, no bacterial growth was observed in the three-phage treatment at both the 4h (*p* =0.001715) and 24h (*p* =0.000191) time points, indicating a successful suppression of phage-resistant bacterial growth and demonstrating a more pronounced effect compared to the other treatments (**Fig. 5d**). Additionally, phage-resistant mutants’ survival in 75% human serum and heat-inactivated (HI) human serum (which served as a negative control) was also examined (**Fig. 5e-g**). ΦBZRI 2, 5 and 6 exhibited significantly lower bacterial loads at 4h when comparing 75% human serum to HI serum (<0.05), whereas ΦBZRI 7 did not showed a significant reduction (**Fig. 5e-g**). These results suggest that ΦBZRI 2, 5 and 6 are more sensitive to serum. This likely indicates that their phage resistance incurs a cost, potentially through capsule defects or surface alterations that render them more susceptible to complement-mediated killing. The HI serum control further supported this, as all mutants exhibited similar growth to the wild-type, indicating that the reduced survival of ΦBZRI 2, 5 and 6 was not merely due to poor growth.

**Fig. 6:**
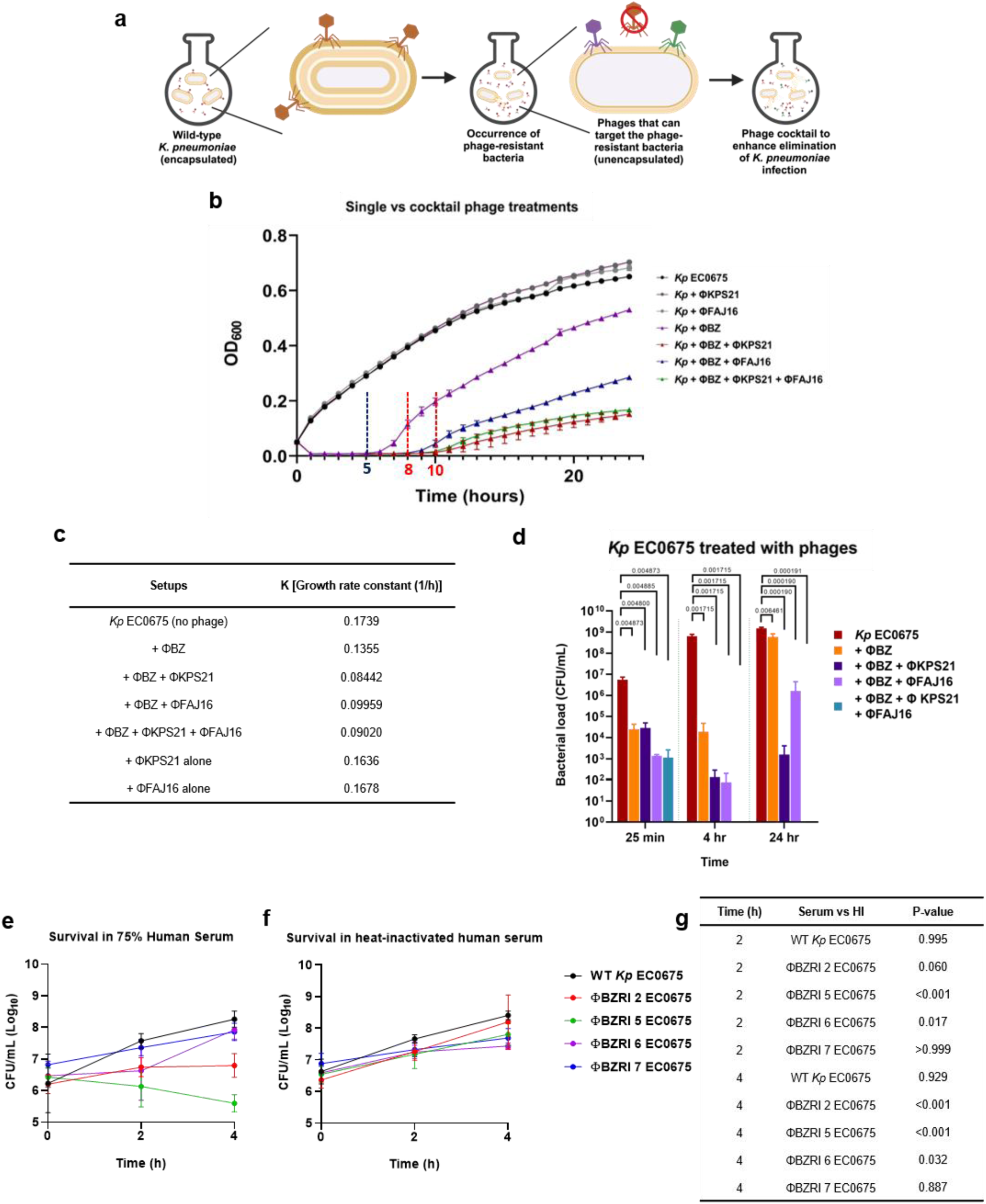
Strategic approach and efficacy assessment of phage cocktail selection. **(a)** Schematic diagram of designing phage cocktail consisting of phages targeting wild-type and phage-resistant populations in the occurrence of phage-resistant bacteria in *kp*. **(b)** The bacterial killing curve of selected cocktail phage treatments against *Kp* EC0675 strain (all data shown at 1h interval readings at OD_600_ represent 3 independent experiments, bars indicate mean ± SEM). **(c)** Growth rate constants (K) of the various phage treatment setups. **(d)** CFU enumeration of single-, two-, and three-phage treatments. n=3 biological replicates; bars indicate mean ± SD. **(e-f)** Bacterial enumeration (CFU/mL) of wild-type *Kp* EC0675 and ΦBZ-RI phage-resistant mutants after incubation in 75% human serum or 75% heat-inactivated human serum for 0, 2, and 4 h. Data are presented as mean ± SD from three biological replicates (n=3). Statistical analysis was performed using two-way ANOVA followed by Šídák’s multiple-comparisons test to compare normal serum with heat-inactivated serum for each strain at 2h and 4 h.

### Assessing phage cocktail against clinical hv-*K. pneumoniae* isolates

The evaluation of the phage cocktail against EC0675 prompted us to validate the combination on a well-characterised collection of 130 *Kp* isolates from a prospective multi-centre randomised clinical trial (Antibiotics for Klebsiella Liver Abscess Syndrome Study, A-KLASS)^46^. *Kp* isolates associated with pyogenic liver abscesses are predominantly hypermucoviscous, hence ideal for our study^32^. Genomic analysis^46^ revealed that A-KLASS strains comprise 68.5% K locus 1 (KL1), ST23, and one KL1 isolate corresponding to ST2044. The non-KL1 isolates include 17% KL2, 4% KL5, 3% KL57, 7.5% other KLs (1.5% KL62/KL124 and 0.78% KL20/KL28/KL63/KL16/KL136/KL127). The sequence type distribution amongst the non-KL1 isolates varies extensively, including ST65, 86, 373, 380, 2039, 420, 20, 592, 36, 111, 399, 60, 828, 1049, 774, 1307, 660, 412, and 2846 (**Extended Data Fig. 3b**). The A-KLASS isolates were categorised into KL1, KL2, and non-KL1/2 heat maps (**Fig. 7a, b**, and **c**, respectively). The data indicate that ΦBZ targets most KL1 (CG23) isolates (n=71/89, 79.8%). In contrast, ΦKPS21 and ΦFAJ16 do not target CG23 isolates, as they were isolated on non-KL1 *Kp* strains, and do not target the wild-type EC0675. ΦBZ did not demonstrate lytic activity amongst the non-KL1 isolates. **Figure 7** highlights a correlation between phage susceptibility and the K-locus in *Kp*. While we have focused on CG23 (KL1), accounting for 67.7% of the isolates, we can see that other phages could potentially be used to target the remaining hv*Kp* isolates associated with pyogenic liver abscess (**Fig. 7**). Our strategy for identifying phages based on clonal group proved successful, as the selected phage ΦBZ targeted most KL1 (CG23) strains. In this case, the clonal group could be correlated to the K-locus and phage susceptibility. Based on these results, we are presently evaluating phages based on the K-locus to target all hv*Kp* isolates, using the A-KLASS isolates as a test model.

**Fig. 7:**
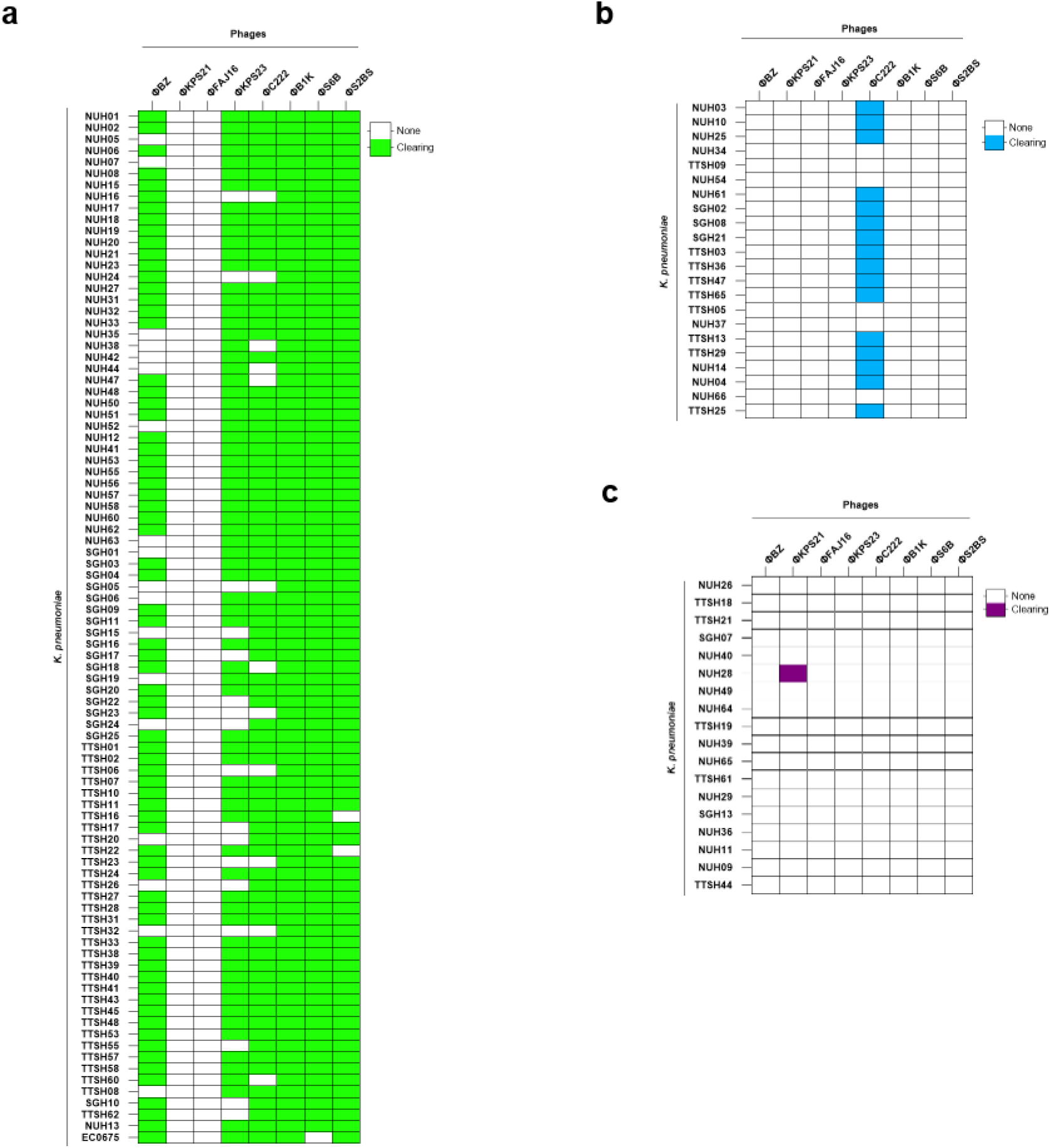
Phage host range profile against 130 *K. pneumoniae* strains in the A-KLASS collection. The phages were spotted on the bacterial overlay of all A-KLASS isolates, and the phage susceptibility profile was tabulated (susceptible: coloured boxes; resistant: clear boxes). **(a)** Phage host susceptibility against CG23 *Kp* strains of capsule serotype KL1. All are ST23 A-KLASS collection except for NUH13 (ST2049). Model strain EC0675 is also included. The last row is *Kp* EC0675 as a comparison with the other CG23 strains. **(b)** Phage host susceptibility against various *Kp* strains of capsule serotype KL2 and mixed STs. **(c)** Phage host susceptibility against various *Kp* strains of other capsule serotypes, non-KL1/2, and mixed STs.

### Phage treatment was effective at clearing *K. pneumoniae* in mice

To evaluate the *in vivo* potential of the selected phages, ΦBZ, ΦKPS21 and ΦFAJ16 were first administered intraperitoneally to mice at 10⁹ PFU to understand the pharmacokinetics and pharmacodynamics and assess their distribution and persistence across organs (**Fig. 8a**). Mice were euthanised at 6h and 24h after phage administration to determine phage load. At 6h, ΦBZ load was observed to be highest in the blood (>7 log_10_ PFU/mL), while the phage load ranged from 5 to 7 log_10_ PFU/mL among the organs **(Fig. 8b)**. At 24h, it was observed that ΦBZ load was significantly reduced in the blood, while the rest of the organs showed no significant reduction. ΦKPS21 and ΦFAJ16 showed lower phage loads (2 to 5 log_10_ PFU/mL) than ΦBZ **(Fig. 8c and 8d)**. It was observed that ΦKPS21 load significantly reduced in the liver and blood while the rest of the organs showed no significant reduction **(Fig. 8c)**. Meanwhile, ΦFAJ16 load showed significant reduction in the lungs and liver while the rest of the organs showed no significant reduction **(Fig. 8d)**. The phage distributions differed only in a few organ pairs, especially with ΦKPS21 and ΦFAJ16 at 6h **(Extended Data Table 2)**. By 24h, broader organ-level significant differences were observed for ΦKPS21 and ΦFAJ16 in comparisons involving the spleen, bladder, kidney and blood. In contrast, both time points showed a more uniform distribution of ΦBZ, with few significant differences in phage load across organs. We next assessed whether phage treatment could reduce bacterial burden in a systemic infection model **(Fig. 8e)**. C57BL/6 mice were intravenously infected with wild-type *Kp* EC0675 and treated intraperitoneally with three doses administered at 3h intervals. Treatment groups received either ΦBZ alone or the three-phage cocktail (ΦCT) comprising ΦBZ, ΦKPS21 and ΦFAJ16. Both ΦBZ and ΦCT were tested at 10⁸ and 10⁹ PFU. At 24h post-infection, all phage-treated groups showed significantly reduced bacterial loads in the liver, lungs, kidneys, spleen, bladder and blood compared with the untreated control **(Fig. 8f)**. The reduction in bacterial burden was more pronounced in the 10⁹ PFU treatment groups, particularly in the liver, lungs, kidneys and spleen, indicating a dose-dependent improvement in therapeutic efficacy. Phage recovery from infected mice confirmed that the administered phages remained detectable at 24h post-infection. No major differences in total phage loads were observed between ΦBZ and ΦCT treatment groups at the same dose **(Fig. 8g)**. Significance was observed between ΦBZ groups and, similarly, between ΦCT groups that showed higher phage load in 10^9^ PFU groups than 10⁸ PFU groups. Within the cocktail-treated groups, ΦBZ was recovered at relatively higher levels than ΦKPS21 and ΦFAJ16 (**Fig. 8g–i**), consistent with the earlier pharmacokinetic analysis. Together, these findings show that intraperitoneal phage treatment can disseminate systemically and significantly reduce bacterial burden across multiple organs in a mouse infection model.

**Fig. 8:**
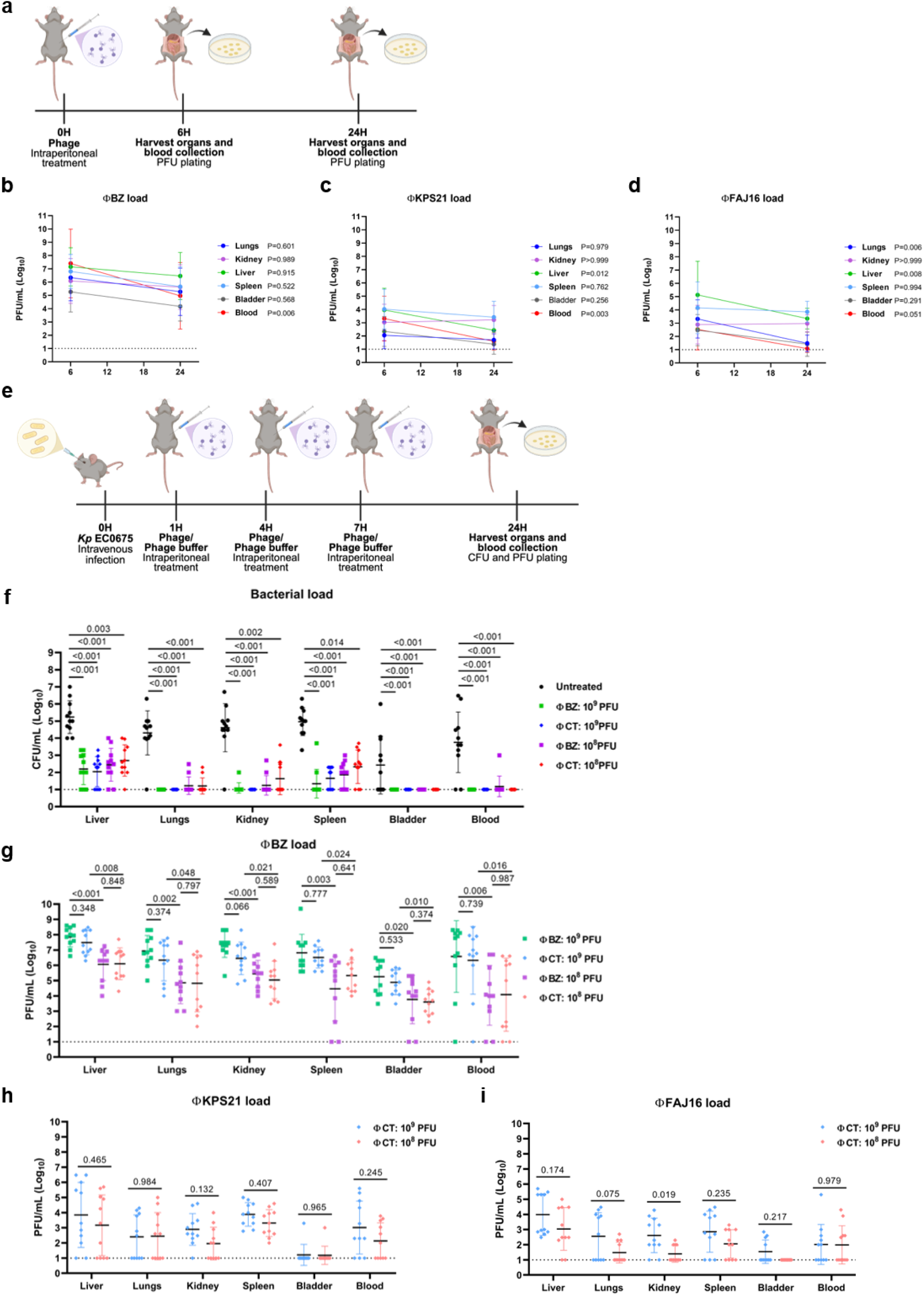
Pharmacokinetics and pharmacodynamics of phages and *in vivo* efficacy assessment of layered phage cocktail. **(a)** Mice experimental timeline of intraperitoneal administration of ΦBZ, ΦKPS21, and ΦFAJ16 at time 0h. Mice were harvested at 6h and 24h for CFU plating. Phage load of **(b)** ΦBZ, **(c)** ΦKPS21, and **(d)** ΦFAJ16 in lungs, kidney, liver, spleen, bladder, and blood following a single treatment administration. Phage titers were determined by plaque assay and expressed as PFU/mL (log_10_) per organ at 6h and 24h post-infection. The lower limit of detection (LOD) is indicated by the horizontal dotted line. Each symbol represents the mean with standard deviation (SD) (n=12). Data were analyzed using two-way ANOVA test followed by Sidak’s multiple comparisons test of the time points (not significant: P>0.05). **(e)** Mice experimental timeline of intravenous infection of wild-type *Kp* EC0675 at 0h, followed by phage treatment at 1, 4, 7h post-infection (hpi). Mice were harvested at 24 hpi for CFU and PFU plating. **(f)** Bacterial load in the organs and blood of untreated and phage-treated mice. Following infection with *Kp* EC0675, mice were treated with ΦBZ or CT (ΦBZ, ΦKPS21 and ΦFAJ16) at 10^8^ or 10^9^ PFU (n≥5 per group). Bacterial load is shown as CFU/mL (log_10_) per organ. Each data point corresponds to 1 mouse, and data were pooled from 2 independent experiments. Each bar represents the mean with standard deviation (SD) (n=11). The horizontal dotted line denotes the LOD. Data were analyzed using Kruskal–Wallis test followed by Dunn’s post hoc multiple comparisons test of phage(s) treatment to the control treatment (not significant: P>0.05). Individual phage loads in the organs and blood of **(g)** ΦBZ, **(h)** ΦKPS21, and **(i)** ΦFAJ16 were quantified with their respective host. Phage load is shown as PFU/mL (log10) per organ. Each data point corresponds to 1 mouse, and data were pooled from 2 independent experiments. Each bar represents the mean with standard deviation (SD) (n=11). The horizontal dotted line denotes the detection limit. Data were analyzed using Kruskal–Wallis test followed by Dunn’s post hoc multiple comparisons test of phage(s) treatments.

### Phage resistance comes at the cost of increased drug vulnerability

Next, we evaluate the potential of the layered phage cocktail to enhance bacterial suppression in combination with antibiotics. *Kp* EC0675 was treated with the phages ΦBZ, ΦKPS21, and ΦFAJ16 in combination with fosfomycin, meropenem, polymyxin B, or ciprofloxacin **(Fig. 9)**. In all conditions, the phage cocktail alone initially suppressed bacterial growth. However, resistance and growth were observed after approximately 8-10 hours, indicating the emergence of phage-resistant populations. Antibiotics alone exhibited variable efficacy: fosfomycin and meropenem **(Fig. 9a and 9b)** effectively limited growth to some extent, whereas polymyxin B and ciprofloxacin **(Fig. 9c and 9d)** provided weaker suppression under the conditions tested. In contrast, combining the phage cocktail with antibiotics resulted in a greater growth inhibition than either treatment alone. This effect was most pronounced with fosfomycin, meropenem, and polymyxin B, where bacterial growth remained significantly suppressed up to 24h. Though the ciprofloxacin combination also reduced growth relative to ciprofloxacin alone, bacterial growth was still observed. Collectively, these findings suggest that phage pressure can lead to phage resistance through the loss of capsules, which in turn unmasks the antibiotic vulnerabilities of drug-resistant bacteria. The layered phage cocktail can augment antibiotic-mediated control of *Kp*, especially when antibiotic treatment alone proves inadequate.

**Fig. 9:**
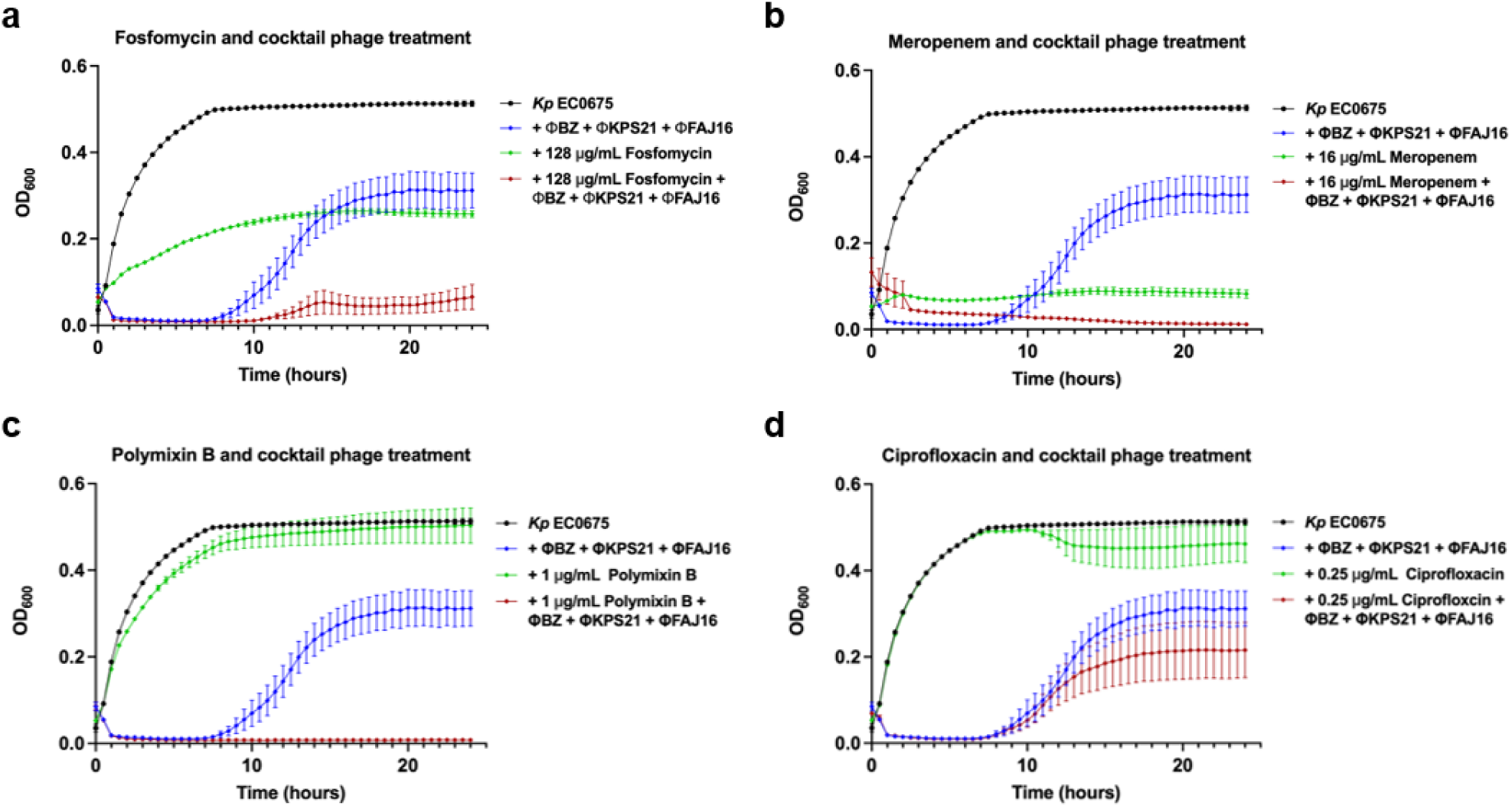
Efficacy of antibiotic-phage cocktail combination treatment. The bacterial killing curve of various antibiotics (a) fosfomycin, (b) meropenem, (c), polymyxin B, (d) ciprofloxacin, with and without the combination of the layered phage cocktail against *Kp* EC0675 strain (all data shown at 30 min interval readings at OD_600_ represent 3 independent experiments, bars indicate mean ± SEM).

## Discussion

Despite promising results^47^, bacteriophages as an alternative therapy to combat multidrug-resistant infections are still hampered by the slow turnaround time to source and characterize suitable phages and the emergence of phage resistance. Using hv*Kp* EC0675, our work addresses both challenges by developing a ready-to-use phage cocktail targeting the regionally important and largest clonal group of hv*Kp* isolates (CG23)^48,49^ associated with as much as 80% of community-acquired pyogenic liver abscesses^46,50,51^ and its alarming trajectory of carbapenem resistance, particularly among ST23 strains now disseminating from Asia across Europe and North America ^52–55^. Its enhanced virulence^17^, combined with rising antibiotic resistance, demands urgent attention as exemplified by CG23 sequence type ST23.

One strategy is to use phage combinations to mitigate or delay phage resistance and improve efficacy. Previous studies have shown that isolating phages based on phage-resistant mutants may provide a rationale for mitigating phage resistance arising during therapy^18,19^. Our results demonstrate that random phage combinations based on genomic or morphological differences showed no synergistic or additive effects. We attribute this lack of effectiveness to shared resistance mechanisms (**Fig. 1d and 1e**). Combining phages based on receptor diversity may not necessarily result in effective mitigation of phage resistance^56,57^. When the bacterium loses its capsule, it may simultaneously lose the receptor surface shared by all CG23-targeting phages, rendering the entire first-layer arsenal ineffective in a single mutational step (**Fig. 2c).** This global cross-resistance, driven by convergent mutations in capsular biosynthesis genes, is also demonstrated in K54 hv*Kp*^58^ is the key reason why conventional diversity-based cocktails fail against this pathogen. Our findings underscore the importance of a strategic “layered” approach^59^ in which the first phage targets the encapsulated wild-type population, while secondary phages are selected against the capsule-impaired mutants that arise during treatment. This design anticipates the dominant resistance pathway and redirects it into a vulnerability that can be exploited by additional phages and antibiotics, thereby improving bacterial suppression and treatment outcomes against hv*Kp* (**Fig. 6a**).

While investigating optimal cocktail design, we found that hv*Kp,* rather than employing diverse resistance pathways, employs CPS loss or disruption driven by point mutations in genes (*wzb*, *wzc*, and *wcaJ*) involved in the biosynthetic pathway, which emerged as the standard mechanism of phage escape^58,60,61^ (**Fig. 3a**). Phage pairing strategies are therefore pathogen-specific; what works against c*Kp* may be actively counterproductive against these hv*Kp* strains. This turned the resistance event into a recruitment filter: only phages capable of infecting the membrane-exposed mutant phenotype were selected for the second layer (**Fig. 4a**).

ΦFAJ16 and ΦKPS21, isolated through this “layered approach”, may target receptors inaccessible on wild-type bacteria. The resulting three-phage cocktail thus operates on two distinct bacterial phenotypes: ΦBZ hits the encapsulated wild-type, while ΦFAJ16 and ΦKPS21 target the capsule-deficient escape mutants that inevitably arise. This strategy delayed the emergence of resistance compared to single-phage treatment, leading to delayed growth amongst resistant population^57^(**Fig. 6b**), suggesting fitness costs due to resistance may translate to therapeutic advantage *in vivo*^62^. Subsequently, the capsule-impaired mutants displayed increased susceptibility to serum killing (**Fig. 6e-g**). The CPS present in encapsulated *Kp* can block the assembly and deposition of complement system components on the microbial surface, thereby inhibiting Membrane Attack Complex(MAC)-mediated lysis^63^. Consequently, the mutants exhibit susceptibility to serum killing but are not completely eradicated. Interestingly, serum sensitivity may be mutation-specific. While the *wzb* and *wzc* mutants showed increased susceptibility to serum killing, the *wcaJ* mutants (ΦBZRI 6 and 7) are more serum-resistant despite impaired capsule production. This may be due to a higher CPS level in the *wcaJ* mutants than in other mutants, as shown in the uronic acid assay (**Fig. 3d**). This is also consistent with previous findings showing that *wcaJ* loss-of-function in *Kp* can reduce capsule production while promoting complement resistance, even with increased complement binding and opsonophagocytosis^64^. Thus, different capsule biosynthesis mutations may reshape host–pathogen interactions in distinct ways, rather than uniformly reducing serum resistance. Altogether, the attenuation of pathogenicity and virulence in capsule-deficient mutants that emerge during phage resistance may provide a critical safeguard against overwhelming infection. Notably, our serum killing assays used pooled healthy-donor serum, and thus may not fully reflect complement and antibody conditions in hv*Kp* liver abscess patients.

The phages in this study were also tested on a representative collection of *Kp* isolates from the pyogenic liver abscess patients from Singapore (A-KLASS study) to validate our findings across a larger population (**Fig. 7**). Against the A-KLASS collection, ΦBZ and 5 additional phages collectively covered all KL1 clinical isolates (**Fig. 7a**). ΦKPS21 and ΦFAJ16 are inactive against encapsulated wild-type KL1, suggesting of limited utility if any; however, their purpose is to intercept the resistant subpopulation that ΦBZ inevitably selects for. This distinction between phages selected for primary activity versus those selected for resistance interception should guide pre-formulation testing in any future clinical cocktail development effort for hv*Kp*.

A key unknown in translating phage cocktails to the clinic is whether second-layer phages administered upfront but not bactericidal against the initial wild-type population can persist long enough to intercept capsule-deficient escape mutants as they emerge. Our PK/PD data in uninfected mice partly address this, as all three phages were recoverable in blood and multiple organs after intraperitoneal injection, confirming systemic dissemination. ΦBZ achieved higher and more sustained titers than ΦKPS21 and ΦFAJ16 (**Fig. 8b–d**), which may reflect phage-specific differences in serum stability or immune clearance. The lower *in vivo* recovery of the second-layer phages is not necessarily a clinical obstacle as they only need to be present at sufficient levels when the resistant subpopulation expands. Because these *in vitro* assays were performed at an MOI of 10, they represent a laboratory stress-test rather than a direct clinical dosing scenario. Effective phage:bacteria ratios at infection sites are often lower, and future work should examine whether the layered cocktail maintains its advantage at lower MOI and in more complex matrices such as biofilms and body fluids. Nevertheless, the fact that capsule-loss remains the dominant resistance pathway under high phage pressure supports its role as a predictable evolutionary escape route in CG23 hv*Kp*. In the systemic infection model, both ΦBZ alone and the layered phage cocktail significantly reduced bacterial burdens in the liver, lungs, kidneys, spleen, bladder, and blood at 24 hours post-infection. This reduction was generally more pronounced at higher phage doses, indicating a dose-dependent therapeutic effect. These findings provide evidence that intraperitoneal phage administration can effectively reduce disseminated hv*Kp* infections *in vivo* and serve as proof-of-concept that a pre-selected phage formulation can target CG23 hv*Kp*. ΦBZ was recovered at higher concentrations than ΦKPS21 and ΦFAJ16, because likely that the initial reduction in bacterial burden was primarily due to the activity of ΦBZ against the encapsulated wild-type population. Meanwhile, the layered components may play a critical role in limiting the expansion of resistant subpopulations over time. In the mouse systemic infection model, both single-phage and cocktail treatments reduced bacterial burden across all six organs at 24h, with consistently greater reductions at 10⁹ PFU than at 10⁸ PFU (**Fig. 8f**). Organ-to-organ variation in efficacy likely reflects differences in tissue perfusion, local bacterial density, and phage persistence. These are factors that will need to be addressed through route and dosing optimisation in future models. The key preclinical question for the layered cocktail is whether the second-layer phages contribute measurably over and above ΦBZ alone at early time points has yet to be addressed, and survival studies will be needed to resolve it. The differential response to antibiotics in the combination experiments is mechanistically coherent (**Fig. 9**). Fosfomycin, meropenem, and polymyxin B, all act on the bacterial envelope or cell wall structures that become directly more accessible when the capsule is lost. Ciprofloxacin targets intracellular gyrase and topoisomerase IV, which are unaffected by surface remodelling; the modest improvement seen with ciprofloxacin likely reflects non-specific population suppression rather than a capsule-loss-mediated mechanism. These observations are consistent with studies showing that phage-driven capsule loss preferentially resensitizes *Kp* to envelope-active antibiotics^65,66^. Together, the data suggest that antibiotic selection for combination regimens should be informed by the predicted mechanism of resistance, rather than chosen empirically.

This study presents notable limitations. Firstly, ΦKPS21 carries genomic features associated with temperate phages (integrase, repressor), which would preclude its use in a clinical formulation without further engineering. However, its inclusion in this study was intentional and serves a specific mechanistic purpose to demonstrate that layer-2 phages targeting capsule-deficient mutants can be identified and that they are active against non-overlapping receptors (TonB-dependent transporter, ferrichrome outer membrane receptor) relative to the first-layer phage. The layered cocktail principle demonstrated here is independent of whether ΦKPS21 itself is deployed clinically, a fully lytic engineered variant, or indeed any phage active against capsule-impaired mutants, would serve the same strategic role. The identification of ΦFAJ16, a confirmed lytic phage, as an equally effective second-layer component underscores this point. Additionally, we are actively searching for alternative capsule-deficient phages for strategic phage pairing. Importantly, resistance ultimately emerged even against the three-phage cocktail, albeit more slowly, and the target bacteria were associated with a loss of fitness. The pairing of two phages in the second layer may prove sufficient, as it is unlikely that resistance will occur simultaneously at two different phage receptors. While a third-layered approach could be explored, it might not be necessary considering the associated loss of fitness and the combination with standard-of-care antibiotic treatment. Phage-antibiotic synergy has been demonstrated to reduce phage resistance and enhance its efficacy in treating the pathogen^67–69^. Moreover, TonB-outer membrane-dependent receptors are involved in various substrate-specific transport processes in Gram-negative bacteria and are associated with phage binding in *Escherichia coli* and *Salmonella*^70–72^, as well as bidirectional phage-antibiotic susceptibility in *Kp*^73^. Finally, this study explored the efficacy of the proposed phage cocktail in mice, but future research focused on developing a consistent animal model for A-KLASS *Kp* strains will enhance testing of strategically designed phage cocktails with antibiotics, allowing for better assessment of clinical translatability. The findings from both animal models and antibiotic analyses highlight the clinical significance of this layered approach. In living organisms, the phage cocktail strategy led to systemic phage exposure and reduced bacterial levels across multiple infection sites. For *in vitro* settings, using the phage cocktail alongside antibiotics resulted in more significant bacterial suppression than when either treatment alone. This indicates that carefully designed layered phage cocktails that account for the possibility of phage resistance could be most effective as complementary treatments, thereby enhancing the effectiveness of current antimicrobial therapies rather than serving as independent substitutes for antibiotics. Future studies should explore whether phage-antibiotic synergy can be reproduced in animal infection models, and how dosing regimens can be adjusted to maintain continuous second-layer phage exposure as resistant mutants emerge. The broader implication of this work is that phage resistance in encapsulated *Kp* is not an unpredictable evolutionary outcome; it is a deterministic consequence of the biology. Capsule loss is the path of least resistance for the bacterium, and it is the same path regardless of which phage initiates the pressure. Cocktail designs that ignore this convergence will fail, whereas designs that anticipates it can channel resistance toward a predictable phenotype that is both immune-sensitive and antibiotic-vulnerable. Whether this principle extends beyond CG23 to other encapsulated *Kp* lineages, or to other encapsulated pathogens more broadly, remains to be explored extensively.

## Materials and methods

### Bacterial strains and culture conditions

All the *K. pneumoniae* strains were cultured in Luria-Bertani (LB) broth or agar (Oxoid) at 37°C, and OD was measured at 600 nm using a spectrophotometer (Shimadzu). The following growth conditions were applied for most assays: single colonies of bacteria were selected per replicate and resuspended in 200 µL of LB broth. Cultures were started at OD_600_ ∼ 0.05, grown in snap cap tubes containing 3 mL of LB broth, and incubated at 37°C with shaking at 110rpm under aerobic conditions. All experiments were performed with cultures grown to an OD_600_ of 0.4 (∼1 x 10^8^ CFU/mL) unless otherwise stated.

### Phage isolation and preparation

Phages used in this study were isolated from various sewages in Singapore using various *K. pneumoniae* hosts. The double agar overlay method, consisting of 0.7% top agar in LB broth with bacterial culture, was aptly adapted as described by Russell and Hatfull (2016) on PhagesDB^74^. Phage stocks were prepared using the double agar overlay method to amplify phages, followed by soaking of the plates in phage buffer (40 mM Tris-HCl of pH 7.4, 75 mM NaCl, 10 mM MgSO_4,_ and 1 mM CaCl_2_) for 4h. The phage lysates were collected and filtered using 0.22 µm syringe filters. The titer (PFU/mL) of each phage was determined on its respective hosts to archive the phage library and for downstream experiments.

### Host range determination

Phage lysates were spotted onto different bacterial overlays and incubated at 37°C overnight. Clearings on the phage spots were indicated as positive host targets, while the absence of clearings suggested no infection.

### Selection of phages tested against hv*Kp* EC0675

Phages that commonly target strains of KL1 capsule type were selected for further evaluation against a clinical hv*Kp* strain EC0675^75^, serving as a model for CG23 (ST23/KL1) strains in the present study. Factors such as plaque morphologies, pre-adaptation to EC0675, and transparency (lytic) of their clearing were considered in the selection of phages. Single phages ΦBZ, ΦS6B, ΦKPS23, ΦC222, ΦB1K, and ΦS2BS were tested against EC0675 in the kinetic kill assay. Subsequently, a random phage pairing involving mainly ΦBZ and other phages targeting the EC0675 wild-type to tackle the increased bacterial load observed in the single-phage kinetic kill assay.

### Kinetic kill assay

For all *in vitro* kill-kinetics assays, we used an MOI of 10 (10 phages per bacterium). This MOI is commonly employed in phage characterisation studies to ensure robust detection of differences in killing kinetics and resistance emergence between treatment arms, while remaining within the range used in other *Kp* phage work. Our choice of MOI 10 is consistent with commonly used ranges for in vitro phage characterisation (MOI 0.1–10)^76,77^. Our primary aim is for comparative purposes to assess how layered cocktails perform relative to single-phage and diversity-based cocktails under identical high-pressure conditions rather than to model absolute clinical dosing. Bacteria culture was grown at 37°C for a few hours and was adjusted to 0.4 (1 x 10^8^ CFU/mL) at OD_600_. Phage stocks and bacterial culture were mixed in equal volumes (100 µL) at a multiplicity of infection (MOI) of 10. The cocktail phages were prepared using equal volumes of the respective phages, entailing the same phage titer and a total volume of 1 x 10^7^ PFU/mL in 100 µL. Readings were recorded at OD_600_ and at intervals of 1h until 24h.

### Phage genome extraction, sequencing and analysis

Phage DNA Isolation Kit (Norgen Biotek Cat. 46800) was used for the extraction of phage DNA using high-titer stocks (>1 x 10^8^ PFU/mL). The phage genome was sequenced using the Illumina HiSeq 150-bp paired-end method services provided by BioBasic and Novogene. Fastqc and fastp^78^ were used to improve assembly quality. The reads were then assembled using SPAdes^79^ with default parameters, followed by annotation using RAST (Rapid Annotations using Subsystems Technology)^80^, “Geneious Prime 2026.1.4 (https://www.geneious.com)” and PHASTEST (https://phastest.ca/)^81^. BLAST was further used to annotate proteins termed “hypothetical”. Genes encoding toxins, virulence, or lysogenic factors were screened. Further analysis of the phage genomes was done to identify the most common phage present in the phage database when compared to ΦBZ, ΦFAJ16, and ΦKPS21 using BLAST. SimpleSynteny webserver was used to visualise and compare synteny across the phage genomes^82^.

### Electron microscopy

Phage and phage-bacteria samples were fixed with 2.5 % glutaraldehyde overnight at 4°C. A 10 µL aliquot of the fixed sample was placed on a 200 mesh made of formvar carbon grid and incubated for 2 minutes for phage samples and 30 minutes for phage-bacteria samples. Following incubation, the grid containing the sample was stained with 1% phosphotungstic acid (pH 7) and consecutively the grid was examined under a transmission electron microscope (FEI Tecnai Spirit G2) at 100kV with an Eagle 4k camera.

### Phage-bacterial resistance assay

The assays were performed in snap-cap tubes containing 400 µL of LB broth, 100 µL of bacterial culture, and phages at an MOI of 10. The volumes of phage buffer and phage stocks varied; whereby, 100 µL of each phage was added for either single or cocktail-phage treatments, and phage buffer was added accordingly to make up the total experimental volume to 1 mL. Experimental control was prepared with equal volumes of phage buffer and bacterial culture. The mixtures were incubated at room temperature for 10 minutes, and 4h and 24h at 37°C at 110 rpm. After incubation, 100 µL aliquots were serially diluted and spread onto LB agar plates in triplicate. All spread plates were incubated overnight at 37°C to evaluate colony morphology and isolation of phage-resistant bacteria.

### Isolation of phage-resistant bacteria

The surviving colonies from the phage-bacterial resistance assay were randomly selected and streaked twice to obtain pure colonies. Selected colonies were spotted with the treated phages to confirm phage resistance after overnight incubation at 37°C. Phages were spotted in 3 µL aliquots, and phage-resistant isolates were streaked twice. Glycerol stocks of these phage-resistant mutants were stored at −80°C for further assays. Phage-resistant mutants from different biological replicates were chosen for whole-genome sequencing.

### Bacterial genome extraction, sequencing, and analysis

The One-4-All Genomic DNA Miniprep Kit (BioBasic Cat no.BS88503) was used for the DNA extraction of *Kp* EC0675, and its phage-resistant isolates. Illumina HiSeq 150-bp paired-end method (BioBasic) was used to sequence bacterial genomes. SPAdes and RAST were used. Illumina raw reads were uploaded onto the Bacterial and Viral Bioinformatics Resource Center (BV-BRC^83^) for Comprehensive Genome Analysis under BV-BRC tools and services. The reads were trimmed, assembled, and annotated (Annotation strategy: Bacteria/Archaea, Taxonomy name: *Klebsiella pneumoniae*) using the default parameters. Kleborate software^84^ was used to obtain taxonomy, acquired virulence determinants, virulence score, serotype prediction, antimicrobial resistance (AMR) determinants, and K/O serotypes of *Kp* strains.

### Identification of mutations

Variation analysis (hosted on the BV-BRC) was performed via mapping the reads of the resistant isolates to a reference genome of the wild-type using BWA-mem^85^, followed by variant calling with FreeBayes. The mutations in phage-resistant isolates were identified and verified via a high fraction of the variant read depth among all the reads at the region corresponding to the variant coverage, along with a high-quality score.

### Viscous string test for hypermucoviscosity

Phage-bacterial resistance assays using phage ΦBZ were conducted for eight CG23 *Kp* clinical strains (EC0675, SGH10, SGH09, SGH25, NUH06, NUH38, TTSH27, and TTSH45), and phage-resistant mutants were isolated from individual strains. The hypermucoviscosity phenotype of *Kp* was evaluated using a previously described viscous string test^86^. Briefly, the bacterial lawns of wild-type and phage-resistant mutants were gently lifted with an inoculation loop to observe any mucoviscous string as it was lifted. Bacterial isolates that produced a string ≥5 mm in length were classified as hypermucoviscous–positive, indicating increased capsule production and a hypermucoviscous phenotype. Meanwhile, they were classified as hypermucoviscous–negative with a string <5 mm, indicating decreased capsule production with no hypermucoviscous phenotype.

### Phage adsorption assay

*Kp* EC0675 wild-type and its phage-resistant isolates were grown to OD_600_ ∼0.4. The bacterial cultures were mixed with ΦBZ at an MOI of 0.01 (10^8^ CFU/mL and 10^6^ PFU/mL) in equal volumes of 500 µL of LB broth and phage buffer. The mixtures were incubated at 37°C. At each time point (0, 1, 5, 10 minutes), 100 µL aliquots of the mixtures were transferred into 10 µL of saturated chloroform, mixed, and then centrifuged at 4000x g for 3 minutes. The supernatants were serially diluted in PBS and spotted in duplicates on *Kp* EC0675 WT overlay plates to enumerate free phage particles.

### Quantitative determination of uronic acid

The capsule production was determined via the quantification of the uronic acid content of the bacteria, involving extraction, purification, and quantification of *Kp* extracellular polysaccharides as described previously with slight modifications ^38,87–89^. First, 500 μL of bacterial cultures grown at 37°C in LB broth to OD_600_ ∼1.0 were centrifuged. The supernatant was decanted, and the pellet was resuspended in 500 μL of PBS. LB broth was included as a blank for OD quantification, using the same steps as those for the bacterial cultures, with the initial PBS resuspension. Briefly, the PBS-resuspended cultures were mixed with 100 μL of extraction buffer (100 mM citric acid pH 2.0 and 1% w/v Zwittergent 3–14 detergent) and incubated at 56°C for 30 minutes. The mixtures were centrifuged at 14,000 rpm at 4°C for 5 minutes. 300 μL of supernatant was aliquoted into 1200 μL of 100% absolute ethanol and left for precipitation on ice for 30 minutes. After centrifugation at 14,000 rpm at 4°C for 5 minutes, the pellet of extracted capsules was washed with ice-cold 70% ethanol and centrifuged again. The supernatant was decanted, and another washing was performed. Then, the pellet was air-dried at 60°C for 10 minutes. 100 μL of Milli-Q water was added to resuspend the pellet. On the ice, 600 μL of ice-cold hydrolysis solution (0.0125 M sodium tetraborate decahydrate in pure sulphuric acid) was added to the 100 μL resuspended pellet and vortexed. The mixtures were incubated at 100°C for 7 minutes and immediately placed on ice to cool down. 10 μL of 0.15% (w/v) carbazole was added into the 700 μL mixture and vortexed. The mixtures were incubated at 100°C for 15 minutes and cooled down at room temperature. A standard curve was also included from 100 μg/mL d-glucuronic acid to 3.125 μg/mL with a series of six 2-fold dilutions, and Milli-Q water as blank. The absorbance of the samples and standards was measured at OD_520_, and the readings of the samples were interpolated into the standard curve for quantification using GraphPad Prism 9.0.0. Quantification of the uronic acid content of the samples was finally normalized to OD_600_ 1.0 to determine as μg uronic acid/OD_600_.

### Gene complementation of phage resistant mutants

The *wzb* gene fragment and the arabinose-inducible backbone (pAraB-Apr) were amplified by PCR from the wild-type strain *Kp-EC0675* and from pCasKP-Apr (Addgene no. 117231), respectively. The two fragments were assembled using the NEBuilder® HiFi DNA Assembly Master Mix (Cat. No. M5520AA) following the manufacturer’s protocol to construct the pAraB_wzb plasmid. The recombinant plasmids were chemically transformed into OneShot™ TOP10 Chemically Competent E. coli (Cat. No.C404003) to amplify the plasmids and were subsequently electroporated into competent ΦBZRI isolates with *wzb* mutation (ΦBZRI 3). The ΦBZRI 3 complementation transformants were confirmed by PCR with pAraB screening primers.

### Layered phage cocktail development

ΦBZ and its combinations did not mitigate phage resistance, thus necessitating a re-strategising of the cocktail development towards a layered approach. The layered phage cocktail developed against *Kp* EC0675 consisted of ΦBZ as the primary phage, which effectively targets the wild-type bacteria, and secondary phages, ΦFAJ16 and ΦKPS21, chosen based on their killing activity against the ΦBZ-resistant population. ΦBZ mainly targets CG23 (ST23/KL1) strains, while ΦFAJ16 and ΦKPS21 broadly target the ΦBZ-resistant isolates, as exhibited in the host susceptibility assay, except for the wild-type bacteria. This approach facilitates the selection of phages to target the persistent phage-resistant population.

### Generation of secondary resistance using isolated resistant bacteria

Various *Kp* EC0675 ΦBZ-resistant isolates were grown to an OD_600_ of ∼0.4 (10^8^ CFU/mL) and mixed with either ΦKPS21 or ΦFAJ16 (10^9^ PFU/mL) at equal volumes of 500 µL. Mixtures were incubated at 37°C, and 100 µL aliquots of the cultures were plated on LB agar after 24h. All spread plates were incubated overnight at 37°C. The secondary phage-resistant bacteria were isolated for the respective phages and randomly selected for whole-genome sequencing and mutation analysis.

### Bacteria killing assay

The bacterial killing assay (BKA) was performed as previously described^90^. Phage stocks of ΦBZ, ΦFAJ16, and ΦKPS21, along with bacterial culture of *Kp* EC0675, were serially diluted and aliquoted in equal volumes (100 µL) across a series of descending dilutions. The phage stocks were prepared in 96-well plates prior to the addition of bacterial culture. Upon mixing, 3 µL of the mixture per well was spotted on LB agar plates for 0, 4, and 24h of incubation at 37°C without shaking. All spot plates were incubated at 37°C overnight.

### Determination of growth rate constant (K) of phage treatment curves

OD_600_ values from the kinetic kill curve assay of ΦBZ, KPS21, and FAJ16 were log_10_-transformed to approximate exponential growth kinetics in GraphPad Prism 8 as log_10_(OD_600_) versus time. The log_10_-transformed graph was then fitted using a nonlinear Gompertz growth model^91^ for characterising bacterial growth that enables the estimation of kinetic parameters, including the maximum specific growth rate (μ), lag time (λ), and carrying capacity (A). For each setup, the growth rate was derived from the slope of the fitted Gompertz curve during the exponential phase. Comparisons of the growth rate constant (1/h) between untreated control and phage-treated setups were performed to evaluate the effect of single and cocktail phage treatments on bacterial growth.

### Bacterial survival enumeration assay

Volumes of 100 – 300 µL of 10^9^ PFU/mL stock solutions of ΦBZ, ΦKPS21, and ΦFAJ16 were aliquoted into 14 mL snap cap tubes. The remaining volume was topped up with phage buffer to a total volume of 500 µL (including both phage lysates and buffer). The control tube, which contained no phage, was prepared with 500 µL of phage buffer. The bacterial culture grew to OD_600_: 0.4 (10^8^ CFU/mL) was diluted to 10^6^ CFU/mL, and 500 µL of this diluted culture was aliquoted into the prepared phage tubes to obtain a final concentration of 2 x 10^5^ CFU/mL. Each setup gives a final concentration of 2 x 10^8^ PFU/mL per phage. Five experimental setups (control, ΦBZ, ΦBZ+ΦKPS21, ΦBZ+ΦFAJ16, and ΦBZ+ΦFAJ16+ΦKPS21) were then incubated at 37°C for time points of 25 minutes, 4h, and 24h. At these intervals, 50 µL from each setup was aliquoted into three wells independently, serving as technical replicates for serial dilution. From these dilutions and the neat samples, 25 µL was plated onto quadrant LB agar plates. After incubating the plates at 37°C overnight, CFU counts were performed. The calculated CFUs were then compared to the control at each time point.

### Serum stability assay

Human serum (pooled) [Zenbio, Cat#: HSER-P50ML, Lot#: SER110320] was thawed overnight at 4°C. As cultures were growing at 37°C in LB broth, half of the human serum was heat-inactivated at 56°C for 30 minutes, which served as a control. In a 96-well plate, 150 µL human serum and 150 µL heat-inactivated human serum were prepared in triplicate. Once the bacterial cultures grew turbid, cultures were centrifuged at 13,000 rpm at 4°C for 5 minutes, and the supernatants were discarded while the pellets were resuspended in PBS. OD_600_ was then adjusted to 0.4, and a ten-fold dilution was performed. Then, 50 µL of the bacterial cultures was added to both the serum and the heat-inactivated serum to obtain a final concentration of 2 x 10^5^ CFU in 75% serum. The mixture was then aliquoted into another 96-well plate for quantification at 0h. The mixtures were immediately incubated at 37°C. At each time point (0h, 2h, and 4h), 22 µL of the mixture was transferred into a 96-well plate, and 10 µL was used for ten-fold serial dilution in PBS. They were then spotted onto LB agar plates and incubated overnight at 37°C to enumerate the CFU. The calculated CFUs were then compared to the control.

### Evaluation of phages on A-KLASS study isolates

In this study, we also evaluated the susceptibility of our phages (ΦBZ, ΦS6B, ΦKPS23, ΦC222, ΦB1K, and ΦS2BS) against the 130 *Kp* strains obtained from the A-KLASS group study. In Singapore, an ongoing clinical trial on the Antibiotics for *Klebsiella* Liver Abscess Syndrome Study (A-KLASS) examines the relationship between the clonal group dissemination and virulence of the *Kp* isolates with the demographics and susceptibility factors of patients^46^.

### Preparahtion of phages for study in mice

Phage stocks (ΦBZ, ΦKPS21, and ΦFAJ16) were concentrated and purified using 5x PEG/NaCl (20% Polyethylene Glycol-8000, 2.5 M NaCl), whereby a 1:4 ratio of cold 5x PEG/NaCl to phage stock was mixed in 1.5 mL snap cap tubes and incubated at 4°C overnight. They were centrifuged at 10,000 rpm at 10°C for 5 minutes, and the supernatant was decanted. Another spin was performed, and the remaining supernatant was carefully discarded using a pipette. The pellet was resuspended in phage buffer and left on ice for 1 hour before all the same phages were pooled. Titer using the respective amplification hosts (*Kp* EC0675, K13, and EC0816) was determined to ensure ≥ 10^9^ PFU/mL was obtained.

### Evaluation of the pharmacokinetics and pharmacodynamics of phages in mice

Female C57BL/6 mice (aged 6-7 weeks) were used for the pharmacokinetics and pharmacodynamics (PK/PD) studies of our phages (ΦBZ, ΦKPS21, and ΦFAJ16). Phages were diluted to ∼3.3 x 10^10^ PFU/mL, and all three phages were mixed in equal volumes. Mice were administered 100 µL of the phage mixture to obtain a final concentration of 10^9^ PFU/mL (for all individual phages) at 0h. At 6h and 24h after the phage administration, mice were euthanised, and liver, lungs, spleen, bladder, and kidney were harvested, with blood collected. Organs were homogenised in 1 mL PBS, and tubes were centrifuged at 10,000 rpm for 5 minutes. Supernatant was aliquoted and serially diluted in phage buffer. Phage titers were determined by spotting on their amplification hosts. All procedures involving mice were reviewed and approved by the Institutional Animal Care and Use Committee of Nanyang Technological University (IACUC Protocol No. A23124).

### Evaluation of phage efficacy in mice

*In vivo* efficacy study in mice was evaluated after intravenous infection of C57BL/6 mice with 10^5^ CFU of *Kp* EC0675. Phage treatment (ΦBZ only, and phage cocktail of ΦBZ, ΦKPS21, and ΦFAJ16) of 10^8^ PFU (for all individual phages) and 10^9^ PFU (for all individual phages) was administered intraperitoneally 1h after infection. Another phage treatment was administered similarly, 4h and 7h after the first treatment. At 24h after infection, mice were euthanised, and liver, lungs, spleen, bladder, and kidney were harvested, with blood collected. Organs were homogenised in 1 mL PBS, and tubes were centrifuged at 400 rpm for 5 minutes. Supernatant was aliquoted and serially diluted in PBS. Bacterial load were determined by spotting onto LB agar with 100 ug/mL carbenicillin. For phage load, tubes were centrifuged at 10,000 rpm for 5 minutes. Supernatant was aliquoted into chloroform to a final concentration of 10% and then vortexed. Supernatant was then serially diluted after centrifuging at 10,000 rpm for 5 minutes. Phage titers were determined by spotting on their amplification hosts.

### Antibiotic-phage kinetic kill assay

The bacterial culture was grown at 37°C for a few hours and adjusted to an OD_600_ of 0.4 (10^8^ CFU/mL). Phage stocks and bacterial culture were mixed in equal volumes (100 µL) at a multiplicity of infection (MOI) of 10. Cocktail phages were prepared in equal volumes of 20 µL per phage, entailing the same phage titer of 5 x 10^9^ PFU/mL to obtain a final 10^8^ PFU. 40 µL of the antibiotics were added to setups with combination treatments. The concentrations of the antibiotics (fosfomycin, meropenem, polymyxin B and ciprofloxacin) selected for this assay were determined from the checkerboard assay (**Extended Data Fig. 3**), which tested various antibiotic concentrations based on the CLSI (https://clsi.org/) guideline of the MIC breakpoints. Readings were taken at OD_600_ and at 30-min intervals for 24 hours.

### Graphical illustrations and schematic diagrams

Graphical illustrations and statistical analysis were prepared using GraphPad Prism 9.0.0, GraphPad Software, San Diego, California, USA (www.graphpad.com). *In vitro* experiments were carried out in triplicate with a minimum of two technical replicates per replicate. Schematic diagrams were designed using the biorender application (www.biorender.com).

## Data and availability

WGS data of KpEC0675 (Accession: SAMN38717950), bacterial strains extracted from the A-KLASS cohort study (Accession: SAMN06112188)^39^, phage-resistant mutants (Accession: PRJNA1377710), phages ΦBZ, ΦFAJ16, and ΦKPS21 (Accession: Submission: 3031311), and phages ΦS6B, ΦKPS23, ΦC222, ΦB1K, and ΦS2BS (Accession: Submission:10859901) have been deposited in the NCBI GeneBank database and are publicly available as of the date of publication.

## Acknowledgments

We thank the students from NUS High School of Math and Science on lab attachment programs for their support in phage purification/amplification.

## Author contributions

PB, AN, and JHL designed the study. AN, JHL, LM, AC, CR, Evangeline, KEL, IB, AK, YHG, BK, RH, and PB conducted the experiments and data analysis. AN, LM, JHL, and PB wrote the manuscript with contributions from all authors. All authors approved the paper and gave their consent for publication.

## Funding

The work was supported primarily by NRF-Competitive Research Programme (NRF-CRP26-2021-0006), by the Lee Kong Chian, Nanyang Technological University Start-up Funds and by the A*STAR Infectious Diseases Labs core research grants by Biomedical Research Council (BMRC) of Agency for Science, Technology and Research (A*STAR) and TF IPC Ltd grant titled “Temasek Foundation Infectious Diseases Programme for Surveillance and Diseases X Resilience” and awarded to P. B. JHL was a recipient of the A*STAR Graduate Scholarship, LM and CR are recipient of the Singapore International Graduate Award (SINGA). The funders had no role in the study design, data collection or analysis, decision to publish, or preparation of the manuscript.

## Conflict of Interests

The authors declare that the research was conducted in the absence of any financial or non-financial relationships that could be construed as a potential conflict of interest.

## EXTENDED DATA

**Extended Data Figure 1.**
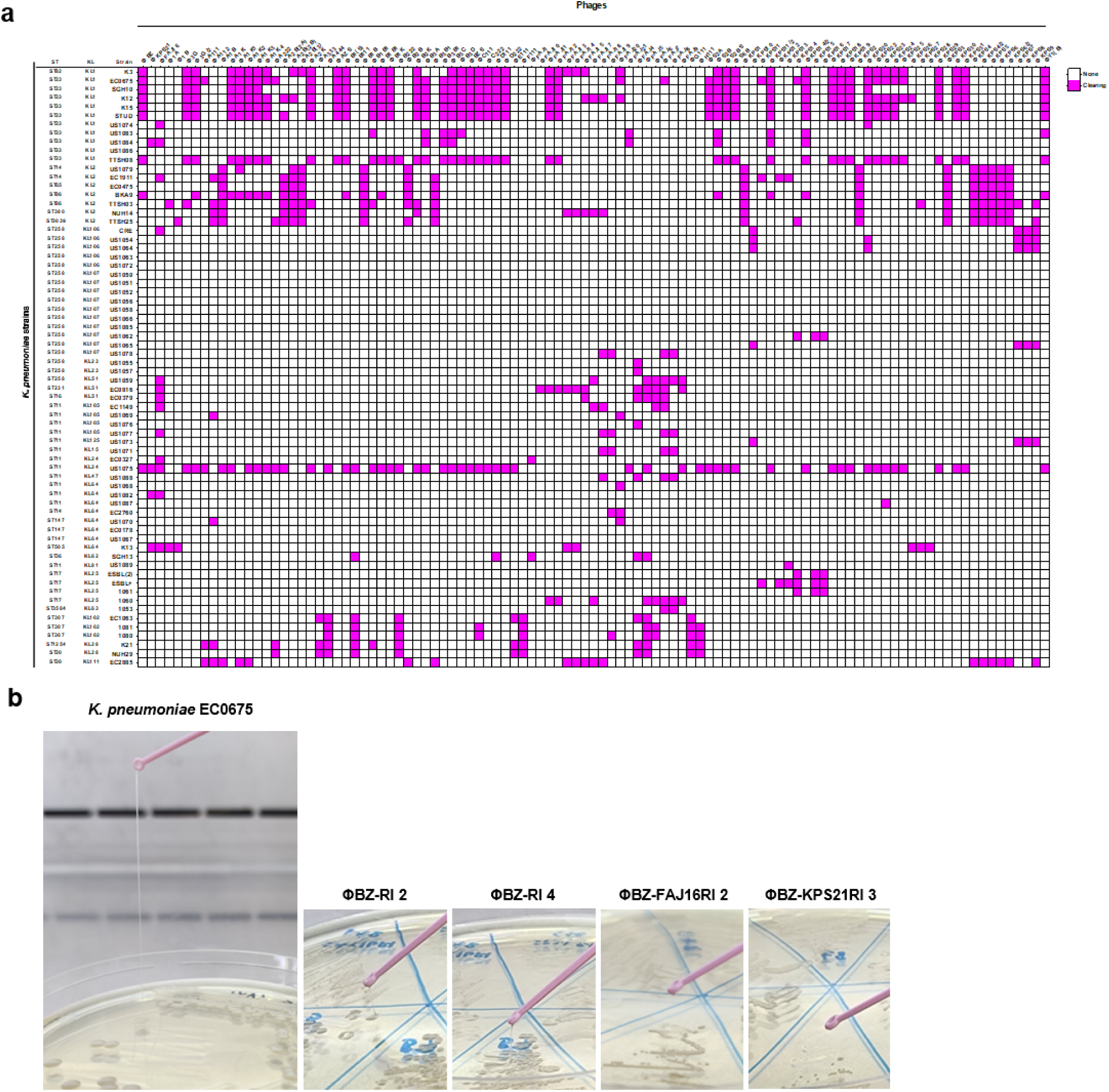
(a) Infectivity profile of phage library against various *Kp* isolates. (b) String test of the bacterial colonies or lawn of *Kp* EC0675 WT and phage-resistant isolates.

**Extended Data Figure 2.**
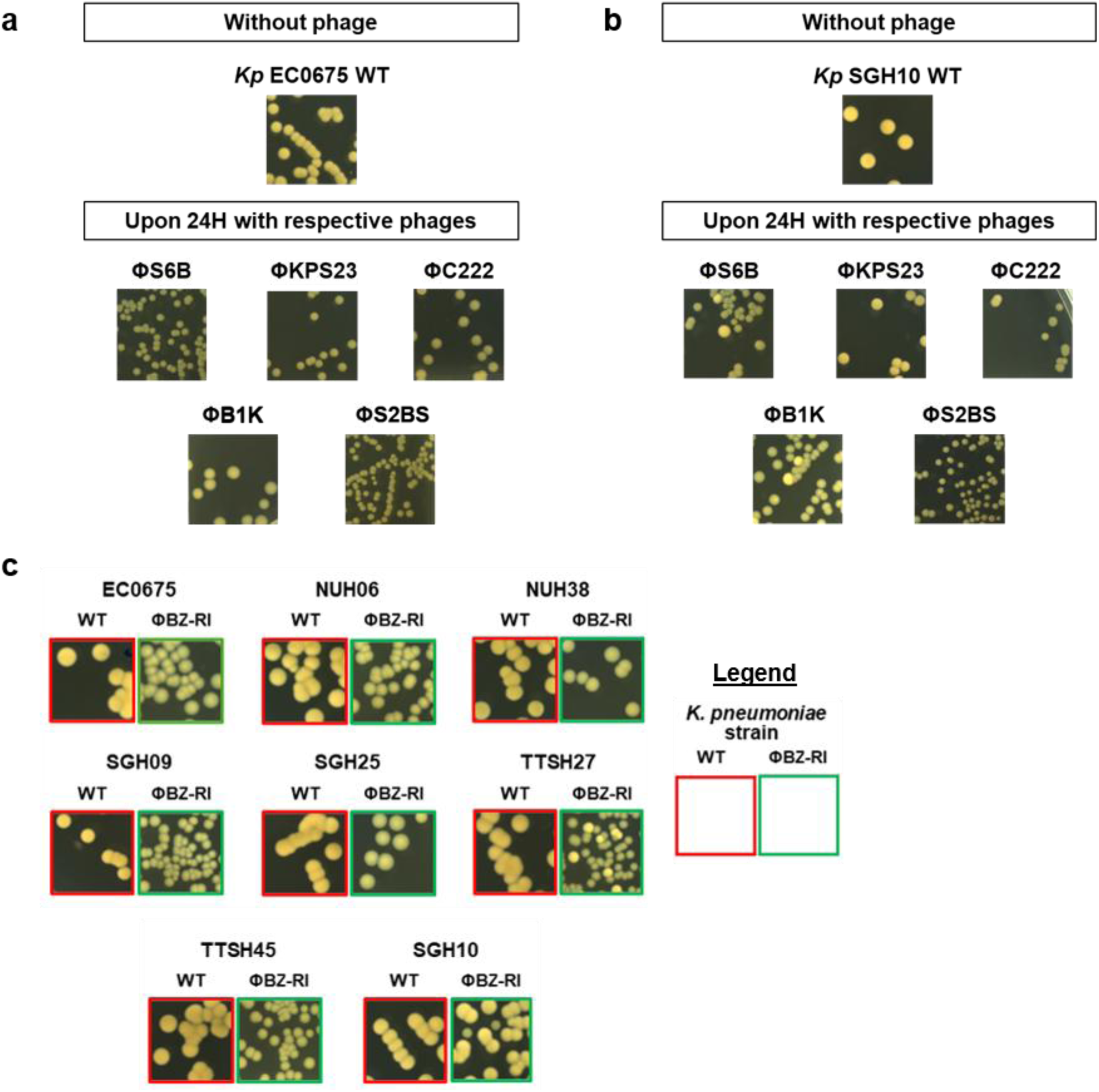
Changes in the bacterial colony appearance of various phage-resistant isolates of **(a)** *Kp* EC0675 and **(b)** *Kp* SGH10 strains in comparison to the wild-type (WT) upon 24h exposure to different CG23-specific phages in liquid *in vitro* infection assay. **(c)** Bacterial colonies phenotype change of ΦBZ-RIs of various CG23 *Kp* strains in comparison to their WT upon 24h exposure to ΦBZ in liquid *in vitro* infection assay.

**Extended Data Figure 3.**
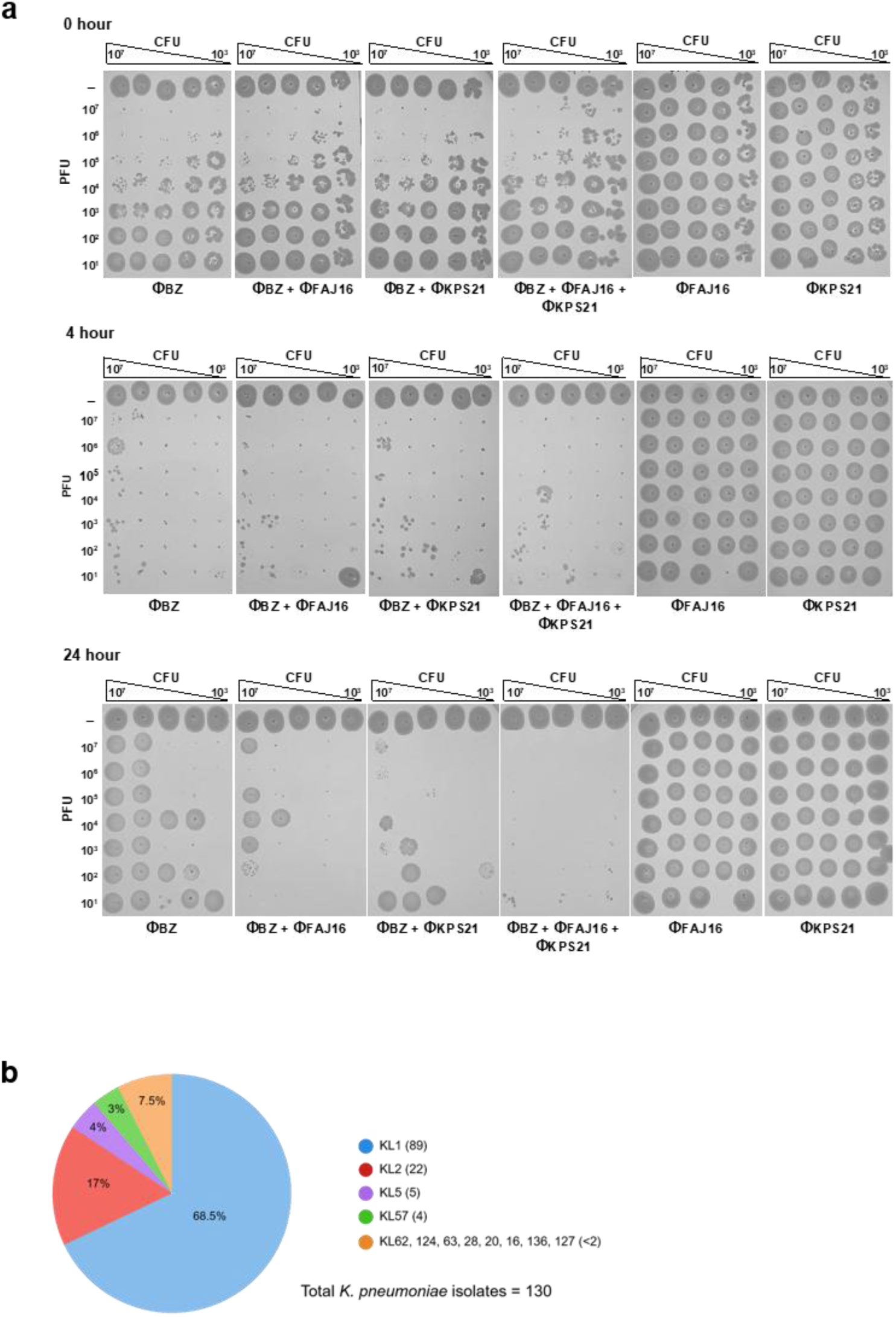
**(a)** The bacterial killing spot assay of single and cocktail phage treatments against *Kp* EC0675 strain (all data shown are representative of 3 independent experiments). **(b)** Proportion of the K locus serotypes present in the *Kp* A-KLASS isolates.

**Extended Data Figure 4.**
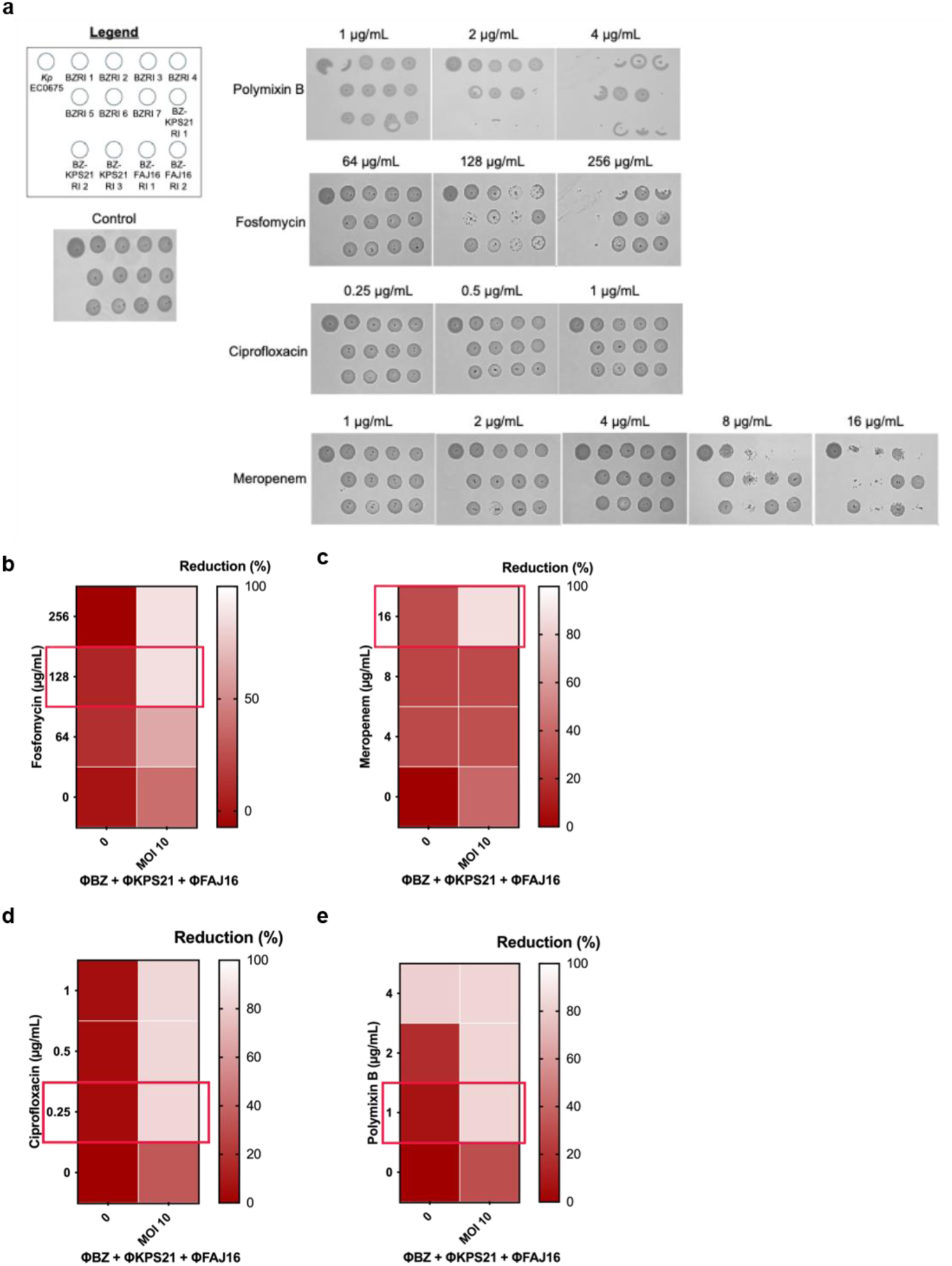
Bacterial growth/reduction of *Kp* EC0675 wild-type and phage-resistant mutants tested on various drugs at various concentrations. **(a)** Bacterial spotting on various drug concentrations and growth after 24h. **(b)** Bacterial reduction (%) of phage cocktail-drug combinations at various concentrations compared to untreated bacteria, measured by OD_600_ after 24h of incubation. Reduction (%) is determined as follows [OD_Kp EC0675 control_ – OD_treatment_/OD_Kp EC0675 control_] x 100%. The concentrations of the antibiotics (fosfomycin, meropenem, polymyxin B and ciprofloxacin) selected for these assays were based on the CLSI (https://clsi.org/) guideline for MIC breakpoints.

**Extended Data Table 1.**
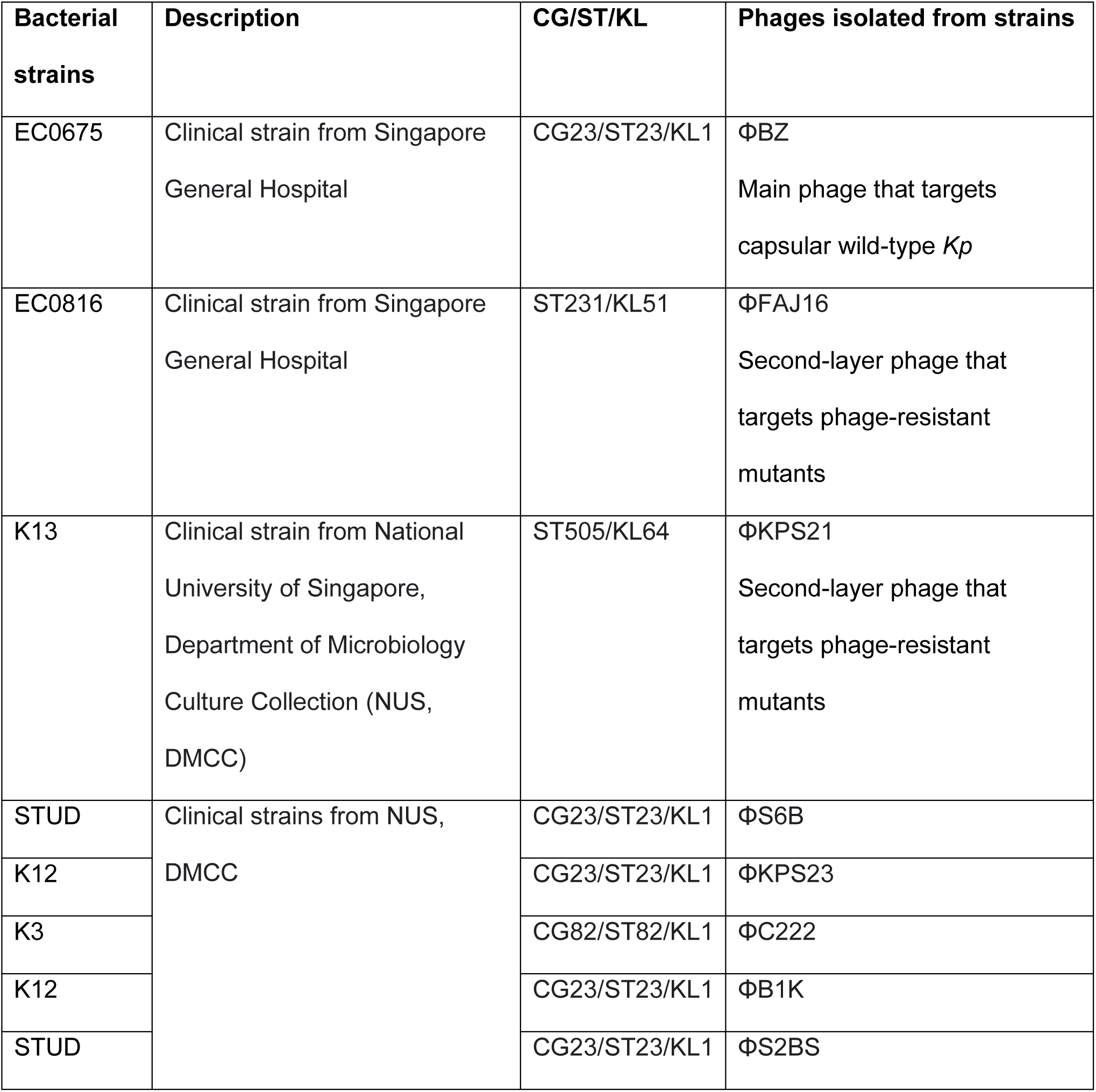
*Klebsiella pneumoniae* strains and phages of interest used in this study.

| Bacterial strains | Description | CG/ST/KL | Phages isolated from strains |
| --- | --- | --- | --- |
| EC0675 | Clinical strain from Singapore General Hospital | CG23/ST23/KL1 | ΦBZ<br>Main phage that targets capsular wild-type *Kp* |
| EC0816 | Clinical strain from Singapore General Hospital | ST231/KL51 | ΦFAJ16<br>Second-layer phage that targets phage-resistant mutants |
| K13 | Clinical strain from National University of Singapore, Department of Microbiology Culture Collection (NUS, DMCC) | ST505/KL64 | ΦKPS21<br>Second-layer phage that targets phage-resistant mutants |
| STUD | Clinical strains from NUS, DMCC | CG23/ST23/KL1 | ΦS6B |
| K12 |  | CG23/ST23/KL1 | ΦKPS23 |
| K3 |  | CG82/ST82/KL1 | ΦC222 |
| K12 |  | CG23/ST23/KL1 | ΦB1K |
| STUD |  | CG23/ST23/KL1 | ΦS2BS |

**Extended Data Table 2.**
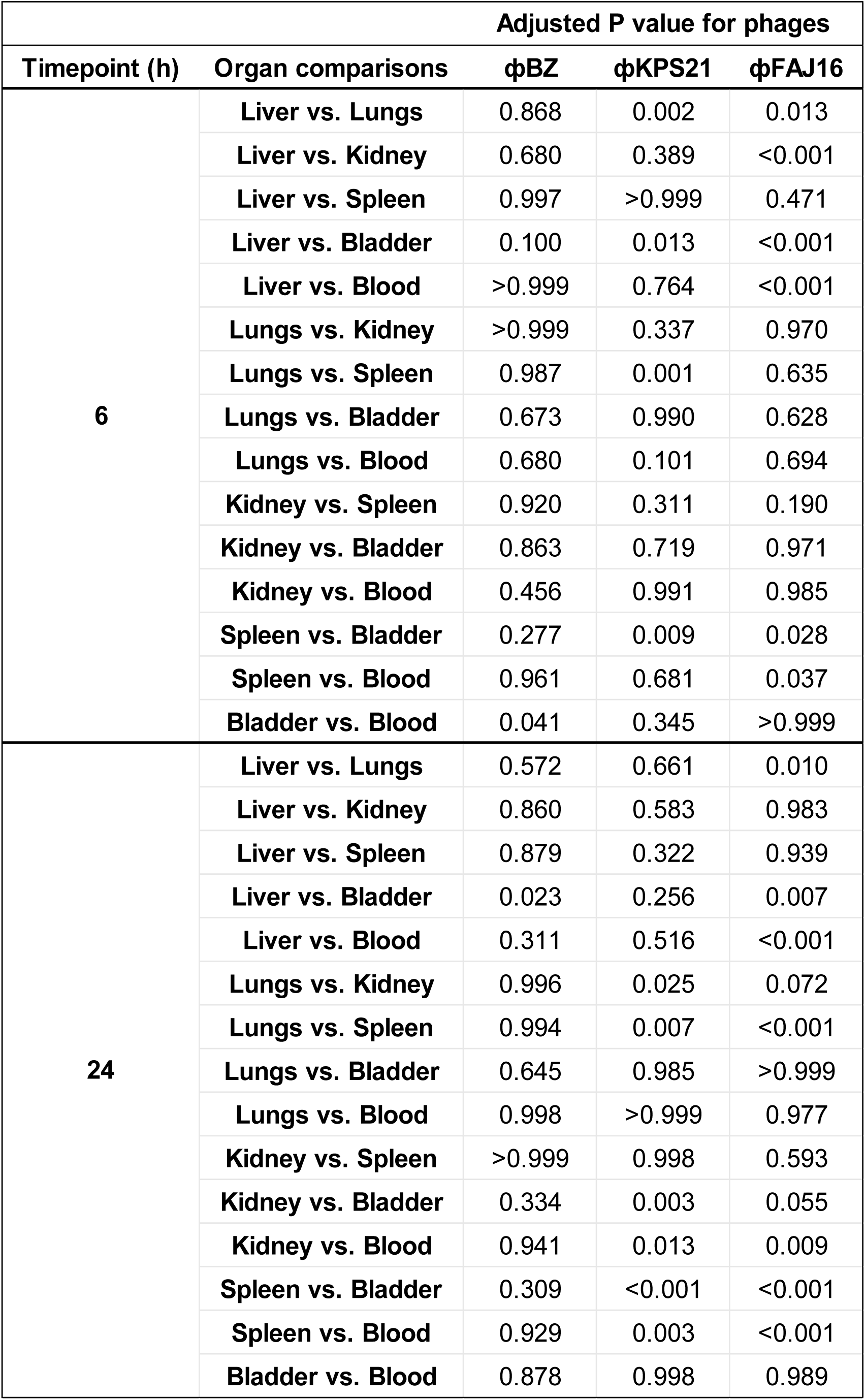
Adjusted p values for pairwise organ comparison of phage distribution at 6 and 24 hours post-administration. Data were analyzed using two-way ANOVA test by Tukey’s multiple comparisons.

## Notes

### Competing Interest Statement

The authors have declared no competing interest.

