## Extended Data Figures for "Layered phage therapy exploits capsule-loss escape and altered antibiotic susceptibility in CG23 hypervirulent *Klebsiella pneumoniae*"

a

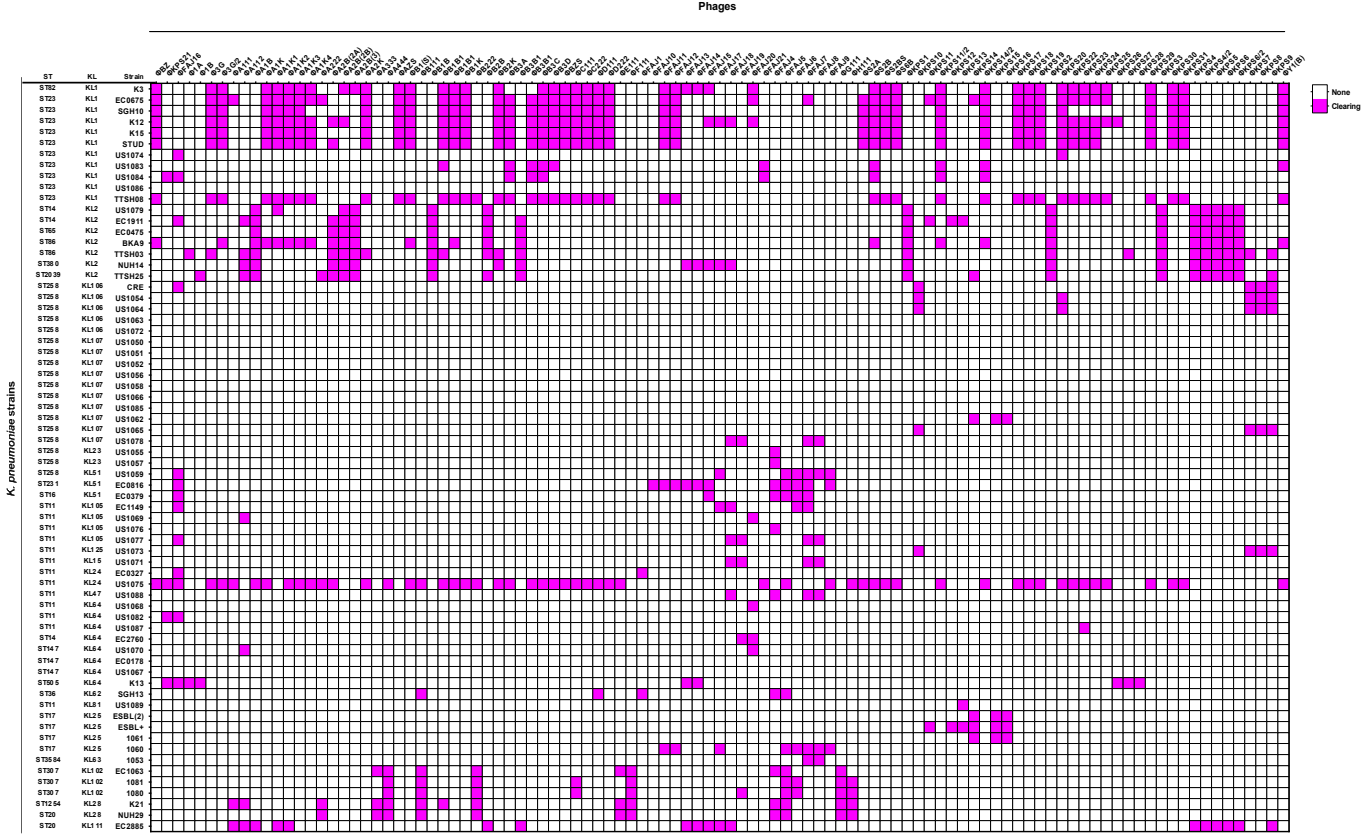

b

*K. pneumoniae* EC0675

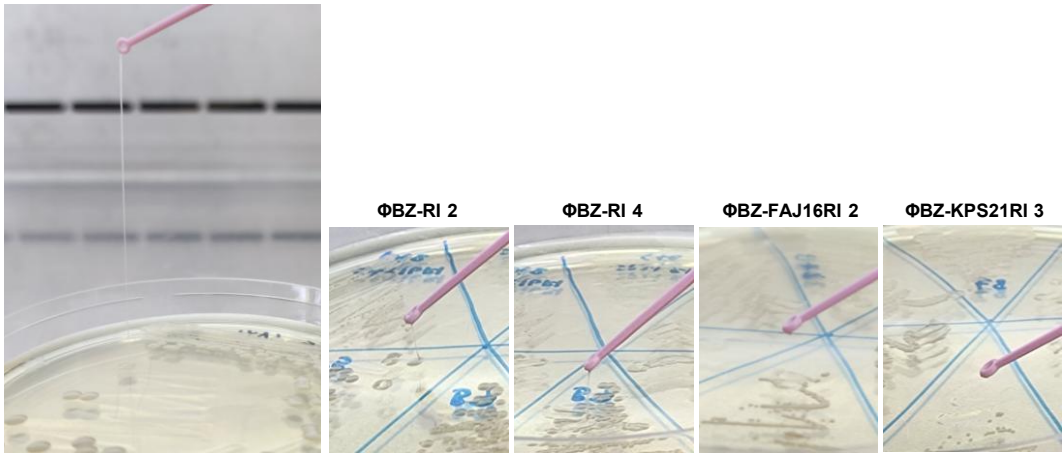

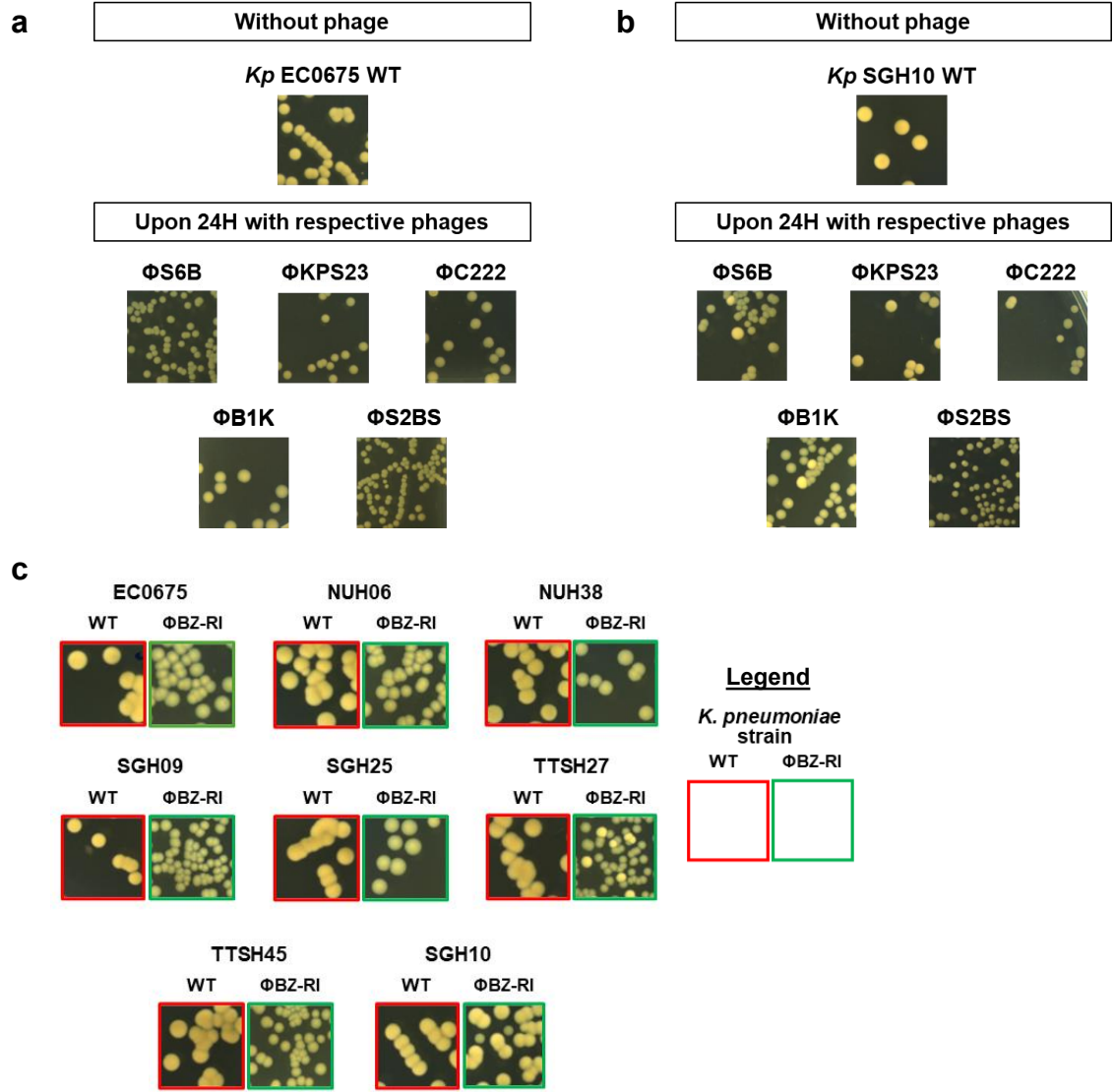

Extended Data Figure 2

a

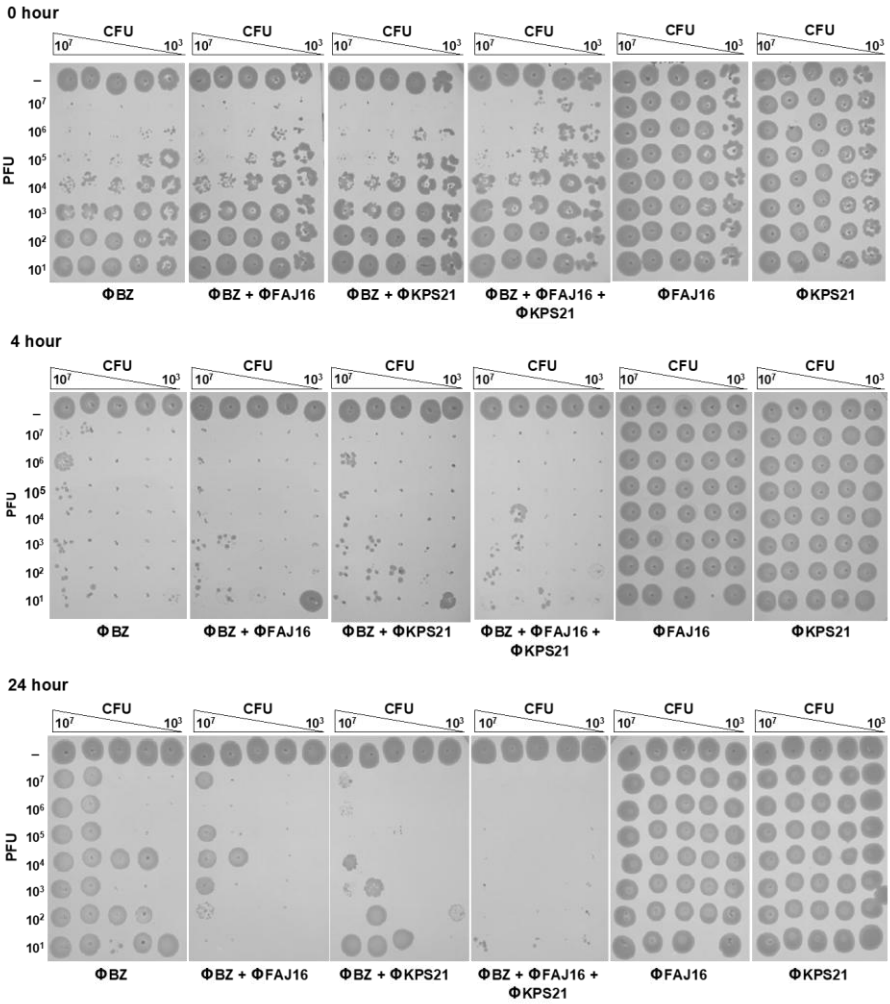

b

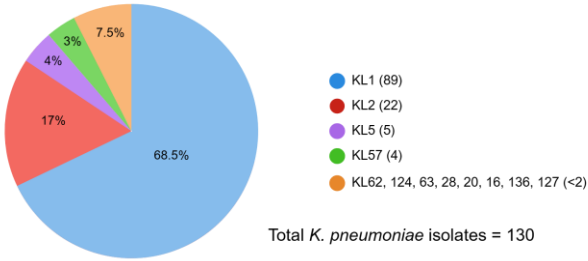

Extended Data Figure 3

**a**

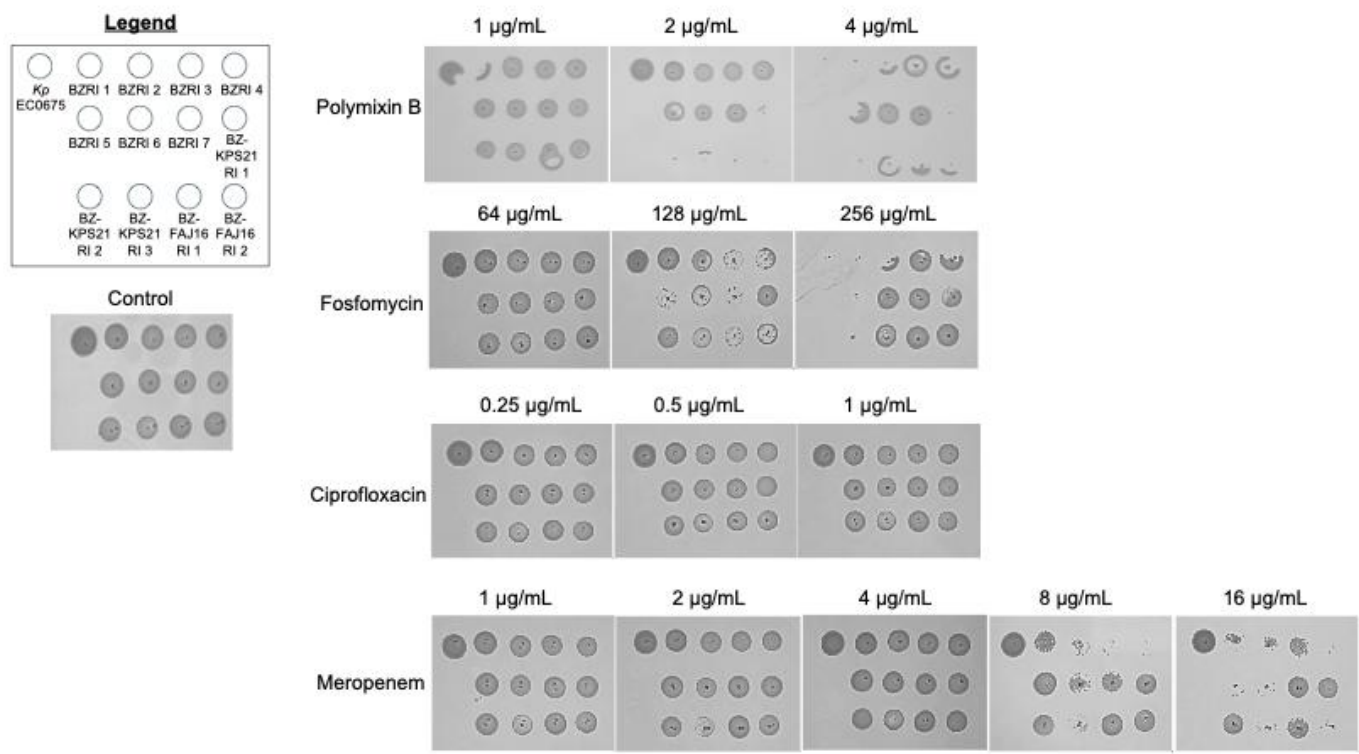

**b**

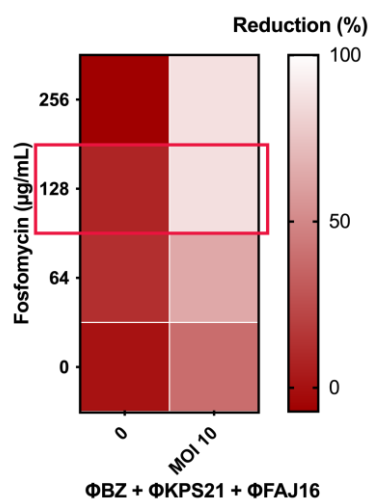

**c**

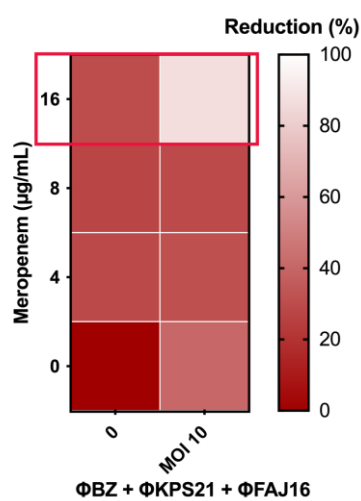

**d**

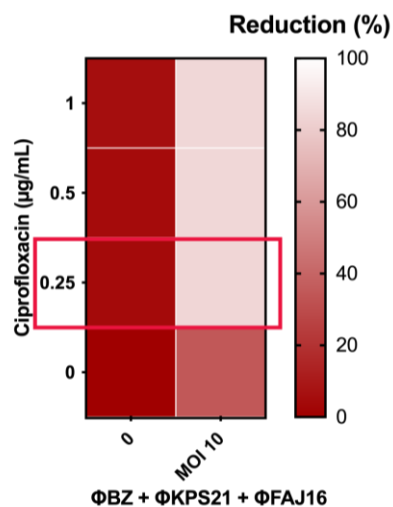

**e**

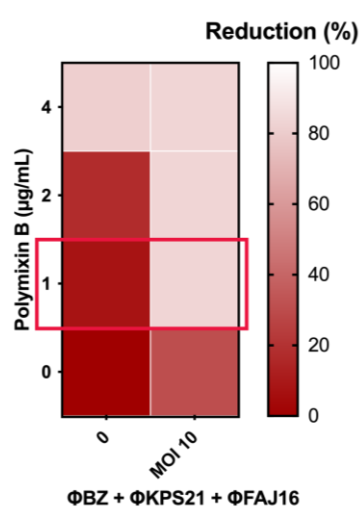

**Extended Data Figure 4**

**Extended Data Table 1.** *Klebsiella pneumoniae* strains and phages of interest used in this study

| Bacterial strains | Description | CG/ST/KL | Phages isolated from strains |
| --- | --- | --- | --- |
| EC0675 | Clinical strain from Singapore General Hospital | CG23/ST23/K<br>L1 | ΦBZ<br>Main phage that targets capsular wild-type *Kp* |
| EC0816 | Clinical strain from Singapore General Hospital | ST231/KL51 | ΦFAJ16<br>Second-layer phage that targets phage-resistant mutants |
| K13 | Clinical strain from National University of Singapore, Department of Microbiology Culture Collection (NUS, DMCC) | ST505/KL64 | ΦKPS21<br>Second-layer phage that targets phage-resistant mutants |
| STUD | Clinical strains from NUS, DMCC | CG23/ST23/K<br>L1 | ΦS6B |
| K12 |  | CG23/ST23/K<br>L1 | ΦKPS23 |
| K3 |  | CG82/ST82/K<br>L1 | ΦC222 |
| K12 |  | CG23/ST23/K<br>L1 | ΦB1K |
| STUD |  | CG23/ST23/K<br>L1 | ΦS2BS |

**Extended Data Table 1**

**Extended Data Table 2.** Adjusted *p* values for pairwise organ comparison of phage distribution at 6 and 24 hours post-administration. Data were analyzed using two-way ANOVA test by Tukey's multiple comparisons.

|  |  | Adjusted P value for phages |  |  |
| --- | --- | --- | --- | --- |
| Timepoint (h) | Organ comparisons | ϕBZ | ϕKPS21 | ϕFAJ16 |
| 6 | Liver vs. Lungs | 0.868 | 0.002 | 0.013 |
|  | Liver vs. Kidney | 0.680 | 0.389 | <0.001 |
|  | Liver vs. Spleen | 0.997 | >0.999 | 0.471 |
|  | Liver vs. Bladder | 0.100 | 0.013 | <0.001 |
|  | Liver vs. Blood | >0.999 | 0.764 | <0.001 |
|  | Lungs vs. Kidney | >0.999 | 0.337 | 0.970 |
|  | Lungs vs. Spleen | 0.987 | 0.001 | 0.635 |
|  | Lungs vs. Bladder | 0.673 | 0.990 | 0.628 |
|  | Lungs vs. Blood | 0.680 | 0.101 | 0.694 |
|  | Kidney vs. Spleen | 0.920 | 0.311 | 0.190 |
|  | Kidney vs. Bladder | 0.863 | 0.719 | 0.971 |
|  | Kidney vs. Blood | 0.456 | 0.991 | 0.985 |
|  | Spleen vs. Bladder | 0.277 | 0.009 | 0.028 |
|  | Spleen vs. Blood | 0.961 | 0.681 | 0.037 |
|  | Bladder vs. Blood | 0.041 | 0.345 | >0.999 |
| 24 | Liver vs. Lungs | 0.572 | 0.661 | 0.010 |
|  | Liver vs. Kidney | 0.860 | 0.583 | 0.983 |
|  | Liver vs. Spleen | 0.879 | 0.322 | 0.939 |
|  | Liver vs. Bladder | 0.023 | 0.256 | 0.007 |
|  | Liver vs. Blood | 0.311 | 0.516 | <0.001 |
|  | Lungs vs. Kidney | 0.996 | 0.025 | 0.072 |
|  | Lungs vs. Spleen | 0.994 | 0.007 | <0.001 |
|  | Lungs vs. Bladder | 0.645 | 0.985 | >0.999 |
|  | Lungs vs. Blood | 0.998 | >0.999 | 0.977 |
|  | Kidney vs. Spleen | >0.999 | 0.998 | 0.593 |
|  | Kidney vs. Bladder | 0.334 | 0.003 | 0.055 |
|  | Kidney vs. Blood | 0.941 | 0.013 | 0.009 |
|  | Spleen vs. Bladder | 0.309 | <0.001 | <0.001 |
|  | Spleen vs. Blood | 0.929 | 0.003 | <0.001 |
|  | Bladder vs. Blood | 0.878 | 0.998 | 0.989 |

**Extended Data Table 2**
